# Dynamin 2–dependent endocytosis differentially regulates ligand and receptor contributions to Notch signaling during neural precursor cell fate determination

**DOI:** 10.64898/2026.06.12.729695

**Authors:** J.J. Breunig, J.I. Arellano, G. Kim, K. Anighoro, C.D. Antonuk, R. Levy, AA Akhtar, J. Molina, M Torii, K. Hashimoto-Torii, S. Fertuzinhos, M.H. Dominguez, L. Grigereit, M. Danielpour, P. Rakic

## Abstract

A critical event in the development of the highly structured cerebral cortex is the appropriate generation of differentiated daughter cell populations from asymmetric precursor cell divisions. Internalization of extracellular receptors and ligands in the process of endocytosis is key to the establishment of asymmetry. However, the detailed mechanisms mediating this exchange are incompletely understood. The dynamin family of membrane remodeling GTPases is considered critical for many forms of clathrin-mediated endocytosis (CME) but its role in brain development is unexplored. Here we found that dynamin 2 (Dnm2), a protein involved in CME through vesicle release from the plasma membrane, is essential for the maintenance of neural stem cells. Conditional deletion of Dnm2 leads to early exhaustion of neural precursor cells and premature neurogenesis, resulting in gross structural abnormalities, periventricular hemorrhaging and increased perinatal mortality. Notably, Notch ligands accumulate at the cell surface and cleaved NICD is reduced in neural stem cells, consistent with impaired ligand-mediated activation. In contrast, inhibition of CME in receptor-expressing cells increases cell autonomous Notch signaling, likely by promoting receptor accumulation at the plasma membrane. Further, dynamin 1, 2, and 3 can rescue loss of Dnm2 to different extents. Taken together, these findings reveal that Dnm2 is the essential isoform for physiological non-cell autonomous Notch-mediated neural stem cell maintenance, and that CME exerts fundamentally distinct roles in Notch signaling, promoting ligand activity while restricting receptor signaling through control of surface receptor abundance.

## INTRODUCTION

Radial glia (RG) are considered the primary precursor population in the developing embryonic forebrain (Noctor, Martinez-Cerdeno et al. 2004, Kriegstein and Alvarez-Buylla 2009, Breunig, Haydar et al. 2011). RG derive from neuroepithelial cells lining the neural tube (Bystron, Blakemore et al. 2008, Breunig, Haydar et al. 2011). During early CNS development these RG divide symmetrically to generate precursors for the pioneering neurons in the brain (Bystron, Blakemore et al. 2008). Later, these RG divide asymmetrically to give rise to intermediate neural progenitors (INPs) that divide symmetrically before migrating into the cortical plate and maturing into functional neurons (Noctor, Martinez-Cerdeno et al. 2004, Breunig, Haydar et al. 2011). During this period, RG provide the substrate for migration and preserve the integrity of the tissue through adhesion at the ventricular (VZ) and the pial surfaces (Rakic 1972, Rasin, Gazula et al. 2007).

Notch signaling has become increasingly appreciated as a central mediator of the cell fate decisions of RG in the developing vertebrate nervous system (Yoon and Gaiano 2005, Ables, Breunig et al. 2011). Notch signaling is one of the most well conserved arbiters of cell fate across metazoa. In the embryonic brain, Notch arbitrates cell-cell communication and in the context of stem cells regulates the choice between self-renewal and differentiation of daughter cells (Yoon and Gaiano 2005, Kriegstein and Alvarez-Buylla 2009). For example, in the nervous system, high Notch signaling is associated with precursor maintenance and low Notch signaling typically leads to differentiation (Breunig, Silbereis et al. 2007, Mizutani, Yoon et al. 2007). Canonically, Notch ligands, such as Dll1, on neighboring cells are thought to stimulate Notch receptors *in trans* by allowing γ-secretase-mediated cleavage of the transmembrane domain of Notch (Kopan and Ilagan 2009), leading to release and nuclear translocation of the intracellular domain of Notch (NICD) and the formation of ternary complex with Rbpj, and MAML-family coactivators (MAML1–3) nuclear factors to initiate the transcription of Notch target genes.

Genetic studies in fly have implicated shibire, the fly homologue of dynamin, as being necessary for effective Notch signaling (Seugnet, Simpson et al. 1997). Subsequently, a host of endocytic molecules, including Neuralized, Mindbomb, and Epsin have been identified as playing a role in the endocytosis of Notch ligands and/or receptors (Fortini and Bilder 2009). For example, in mouse CNS development Mib1 has been shown to be necessary in signal sending cells for the proper endocytosis of Notch ligands during mouse CNS development (Yoon, Koo et al. 2008). However, some endocytosis-related molecules which are critical for invertebrate Notch signaling appear to be dispensable, or are rendered functionally redundant for Notch signal activation in mammals. For example, while in the fly, Mindbomb and Neuralized are necessary for ubiquitation of ligands and activation of Notch in different developmental contexts (Lai and Rubin 2001, Itoh, Kim et al. 2003), in mammals, the two neutralized homologues, and Mib2 appear to be dispensable for development Notch signaling, leaving the aforementioned Mib1 as the critical mediator of ligand ubiquitination (Koo, Yoon et al. 2007, Yoon, Koo et al. 2008).

Despite the growing body of evidence of the necessity of endocytosis for proper ligand function, the necessity of receptor endocytosis for Notch activation remains controversial (Raimondi, Ferguson et al. 2011). In fly, it appears that Notch extracellular domain must be transendocytosed prior to γ-secretase cleavage as dominant negative dynamin blocks the generations of NICD and Notch reporter activity (Vaccari, Lu et al. 2008, Windler and Bilder 2010). Initial studies in mammalian cell lines have also demonstrated the requirement for Notch receptor endocytosis for proper signaling (Gupta-Rossi, Six et al. 2004). On the other hand, some early fly and C. elegans studies indicated that dynamin-dependent internalization of the Notch receptor was dispensable for subsequent receptor activation (Struhl and Adachi 2000, Shaye and Greenwald 2002). Similarly, in mammals several groups have reported that receptor endocytosis is dispensable for Notch cleavage and that most γ-secretase-mediated cleavage happens at the cell surface (Tagami, Okochi et al. 2008, Sorensen and Conner 2010). However, others have pointed out that most, if not all, of these studies rely on overexpression of the full-length receptor or truncated forms, which lack the Notch extracellular domain but require further γ-secretase cleavage for activation (Vaccari, Lu et al. 2008, Sorensen and Conner 2010). Thus, the role of receptor endocytosis in physiological Notch protein cleavage *in vivo* and the potential critical players in this process in vivo remain incompletely understood.

Clathrin mediated endocytosis represents an evolutionarily conserved mechanism for the internalization of plasma membrane proteins(Conner and Schmid 2003). It is used in all cell types for diverse purposes including the regulation of cell surface abundance of receptors and cell adhesion molecules and the uptake of nutrients (Conner and Schmid 2003). A large collection of proteins with roles in cargo recognition, lipid metabolism, membrane deformation and actin cytoskeleton regulation have been identified that make up the machinery that controls clathrin mediated endocytosis (Conner and Schmid 2003). Of these endocytic proteins, the dynamin GTPases are amongst the most investigated and have well characterized actions in the final membrane fission step of this pathway (Ferguson and De Camilli 2012). In addition to the prominent endocytic membrane fission function for dynamins, additional roles in signaling and regulation of the actin cytoskeleton have also been proposed based on studies of dynamin localization and protein-protein interactions(Ferguson and De Camilli 2012). Mammalian genomes encode 3 dynamin genes which share common domain organization but which differ most significantly in their expression patterns (Ferguson and De Camilli 2012). In the nervous system, the embryonic expression pattern of these three molecules is poorly defined and their potential role in development is unexplored. However, in the postnatal brain, Dynamin 1 is most strongly expressed in neurons (Cook, Mesa et al. 1996, Cao, Garcia et al. 1998). A key role for Dynamin 1 in contributing to the efficiency of synaptic vesicle recycling has been substantiated by recent studies in knockout mice (Ferguson, Brasnjo et al. 2007). Dynamin 3 has also been implicated in the function of mature neurons (Raimondi, Ferguson et al. 2011).Dnm2 is ubiquitously expressed and thought to participate in endocytic functions that are common to all cell types (Cook, Mesa et al. 1996, Cao, Garcia et al. 1998, Ferguson and De Camilli 2012). Consistent with such widespread actions, the Dnm2 knockout mouse has an early embryonic lethal phenotype(Ferguson, Raimondi et al. 2009). However, specific functions within the nervous system have neither been identified nor ruled out. Therefore, we have generated CNS-specific conditional mutants for Dnm2 to examine the role of this protein in CNS development and stem cell maintenance.

## RESULTS

### Conditional Inactivation of Dynamin 2 in the Embryonic CNS

We previously generated a floxed Dynamin 2 allele for the examination of the role of Dynamin proteins in the formation of clathrin coated pits (Ferguson, Raimondi et al. 2009). Nestin starts expression around E10.5 (Hockfield and McKay 1985, Frederiksen and McKay 1988) and the Nestin-Cre line of mice starts driving recombination at E11.5 in the CNS (Chou, Perez-Garcia et al. 2009). Nestin-Cre animals were crossed with these conditional Dynamin 2 animals in order to generate CNS-specific Dynamin 2 mutants (Tronche, Kellendonk et al. 1999). Conditional Nestin-Cre Dnm2 mutants (Nes-Dnm2 cKO) died after birth and exhibited a thinned and optically translucent cortex. Examination during embryonic development reveals that up to E12.5, the brain of mutant animals appears indistinguishable from control littermates, but at E13.5, subtle structural changes can be noticed, with the surface of the ventricular zone (VZ) appearing uneven and irregular and by E14.5 tissue lamination is clearly disturbed (**Fig. 1A-F**). To test the efficiency of recombination, we collected forebrain protein lysates at E16.5 from control littermates and Nes-Dnm2 cKO mice. A Dynamin 2-specific antibody indicates an almost complete loss of Dynamin 2 from homozygous animals (**Fig. 1G**). In addition, a pan-Dynamin antibody, which recognizes all isoforms, displayed that Dynamin 2 is the predominant form of Dynamin expressed at this time as we were unable to detect Dynamin protein in the Dynamin 2 mutants (**Fig. 1G**). Together, these data demonstrate that neural progenitor–specific loss of Dynamin 2 disrupts early forebrain architecture and ventricular zone organization, establishing a critical requirement for Dynamin2-mediated membrane trafficking during cortical development.

**Figure 1.**
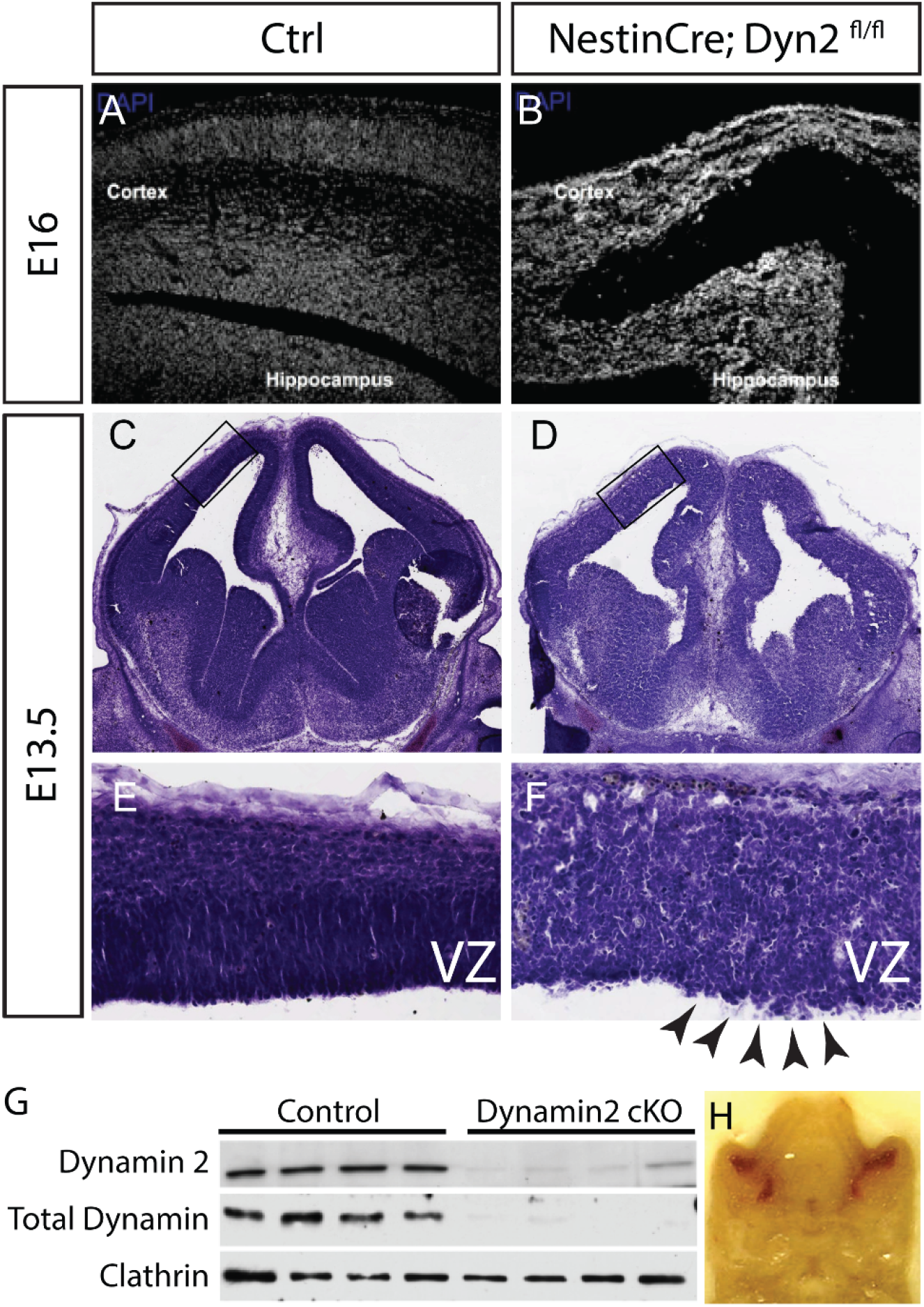
Conditional loss of Dynamin 2 disrupts ventricular zone architecture during forebrain development. (A,B) Representative coronal sections of embryonic cortex from control (Ctrl; A) and Nestin-Cre; Dnm2^fl/fl^ (Dnm2 cKO; B) embryos stained with **DAPI**, illustrating disrupted cortical organization in Dnm2 cKO. (C–F) H&E-stained coronal sections of control and Dnm2 cKO forebrains across embryonic stages, as indicated, showing progressive architectural abnormalities and cortical hypoplasia in Dnm2 cKO. (E,F) Higher-magnification views of dorsal ventricular zone (VZ) regions (boxed in C,D), highlighting early disruption of ventricular surface organization in Dnm2 cKO. (G) Immunoblot analysis of embryonic forebrain lysates from control and Dnm2 cKO littermates showing near-complete loss of Dynamin 2 protein in Dnm2 cKO. A pan-Dynamin antibody reveals that Dynamin 2 is the predominant dynamin isoform in embryonic forebrain at this stage. Clathrin serves as a loading control. (H) Representative image of an embryonic Dnm2 cKO brain showing intraventricular/periventricular hemorrhage.

A detailed analysis with cell markers demonstrated clear changes in the organization of the VZ and SVZ as early as E12.5, whereby Sox2+ progenitors appear detached from the VZ surface and form detached clusters, and the nestin+ radial glia scaffold is disrupted (**Fig. 2**). The cell cycle marker Ki67 shows the characteristic ribbon of intensely stained cells corresponding to dividing progenitors in the VZ surface at E11.5 but disappears by E12.5 in the mutants, with dividing cells distributed throughout the VZ rather than confined to the apical surface (**Fig. 2A-D**). Overall, there is a progressive loss of tissue organization and by E14.5, tissue lamination was noticeably disturbed in all Nes-Dnm2 cKO mice (**Fig. 2E-H**). There was no distinguishable difference at any age between Dnm2fl/fl animals negative for Nestin-Cre and either Dnm2 wt/wt and Dnm2fl/+ animals positive for Nestin-Cre (data not shown). Further, investigation of a comprehensive mouse developmental single-cell RNA sequencing atlas (Qiu, Martin et al. 2024) indicates that Dnm2 mRNA expression is relatively ubiquitous across age, stage, and organ while Dnm1 and Dnm3 are enriched in more differentiated cell types—specifically oligodendrocytes and neurons in the brain (**Fig. S1A-P**). (Notably, though Dnm3 mRNA is enriched in some differentiated intermediate progenitor types as defined by the labeling terminology in this atlas, the precise cells expressing Dnm3 are more akin to differentiating neurons based on their co-expression of synaptic mRNAs as well as Rbfox3, and Neurod2 [**Fig. S1B, E-H**].) These results demonstrate that Dynamin 2 is the principal isoform of Dynamin expressed in the embryonic CNS and that loss of this isoform leads to a hypoplastic neocortex. These findings indicate that Dynamin 2 is required to maintain normal progenitor proliferation and Sox2/Nestin-positive radial glial organization during early corticogenesis.

**Figure 2.**
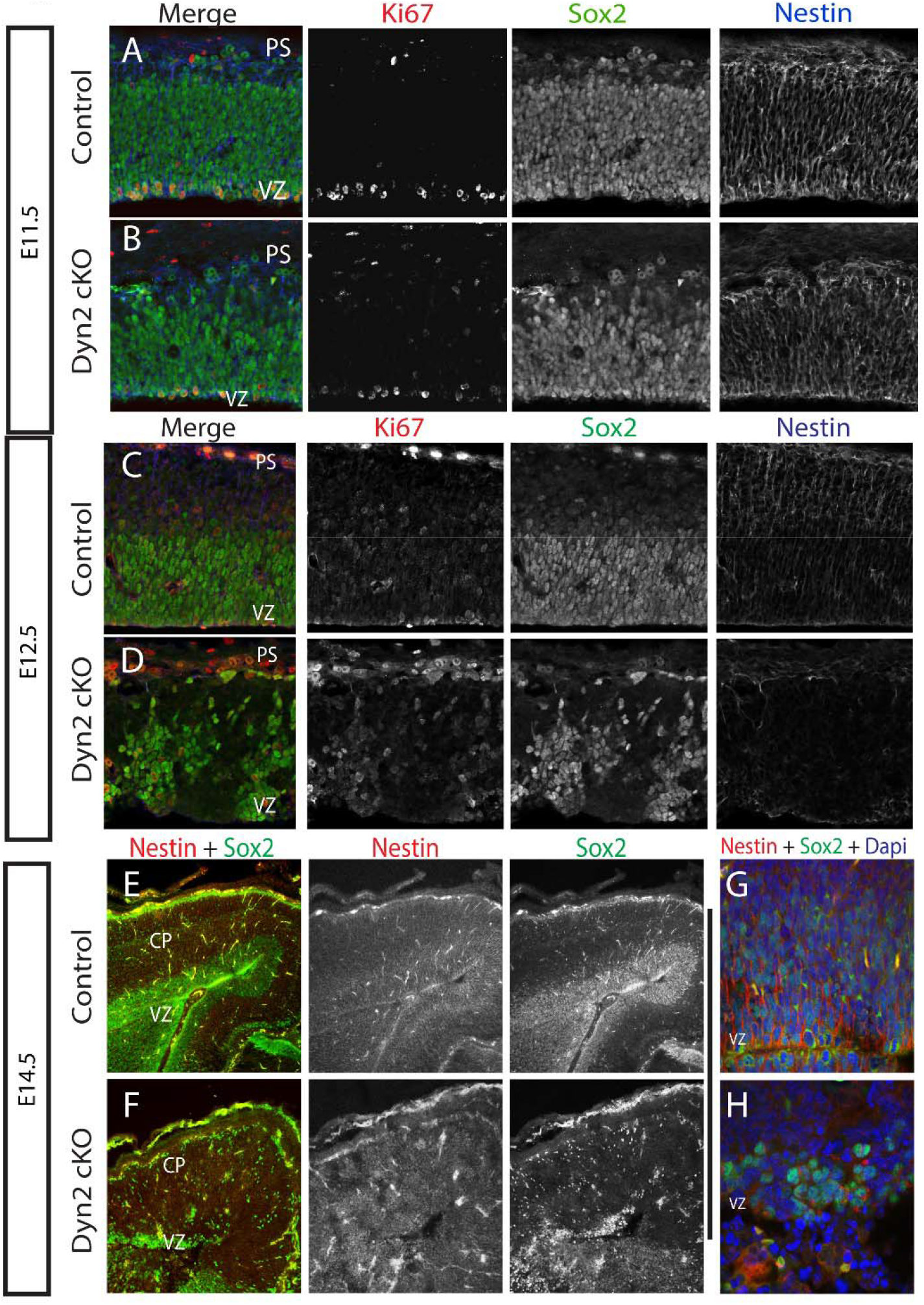
Progressive depletion and disorganization of neural stem and progenitor populations in Dnm2 cKO cortex. (A,B) Confocal images of E11.5 coronal cortical sections from control (A) and Dnm2 cKO (B) embryos immunostained for Ki67 (proliferating cells), Sox2 (neural stem cells), and Nestin (radial glial processes), with DAPI. In control cortex, Sox2^+^ progenitors form a dense apical ventricular zone (VZ), whereas in Dnm2 cKO cortex Ki67^+^ cells are reduced at the ventricular surface and redistributed throughout the VZ, with early disruption of Nestin^+^ radial fibers. (C,D) Confocal images at E12.5 showing further loss of apical organization, with Sox2^+^ cells forming aggregates displaced from the ventricular surface and fragmentation of radial glial processes. (E,F) Higher-magnification views of the VZ at E12.5. Control cortex exhibits a compact, radially aligned progenitor layer, whereas Dnm2 cKO cortex shows thinning, discontinuity, and local breaches of the ventricular surface (arrowheads).

### Lack of Dynamin 2 in the dorsal forebrain produces early depletion of progenitors and increased neurogenesis

Since Nestin is a generic marker of nervous system progenitors, we generated cortex-specific mutants by breeding floxed Dnm2 and Emx1-Cre carrying mice (Iwasato, Nomura et al. 2004). Emx1 is a transcription factor expressed in progenitors and neurons of the dorsal telencephalon that starts expression at E9.5 (Simeone, Acampora et al. 1992). As in Nestin-Cre animals, Emx1-Cre driven recombination is delayed, starting at E10.5 (Chou, Perez-Garcia et al. 2009). At this age, the developing cortex of mutant embryos looks no different than the controls (**Fig. 3**), although close inspection reveals a variable amount of pyknotic cells in the dorsal VZ not present in controls (**Fig. S2**). At E11.5, a drastic phenotype becomes evident, with a disrupted morphology of the dorsal cortex that exhibits loss of normal lamination and a large accumulation of cells protruding into the ventricle (**Fig. 3E**). Notably, electron microscopic analysis of the VZ of E11.5 Emx1-Cre cKO Dnm2 mice demonstrates increased numbers of clathrin coated vesicles and tubular extensions in Dnm2 cKO cells versus control VZ populations (**Fig. 3G-H**) akin phenotypes observed in Dnm2 cKO cells in vitro (Ferguson, Raimondi et al. 2009). This phenotype quickly extends laterally so by E12.5 covers most of the developing cortex (**Fig. 3F**) and by E13.5 the dorsal forebrain is reduced to an atrophied strip of tissue covering the basal forebrain (data not shown). This gradual phenotype seems at odds with the spread of Emx1 expression. However, close inspection of the literature showed that even when there is no explicit mention of an Emx1 early expression gradient, prior studies indicate an early medial-to-lateral expression gradient that would reconcile this observation (Yoshida, Suda et al. 1997, Iwasato, Nomura et al. 2004).

**Figure 3.**
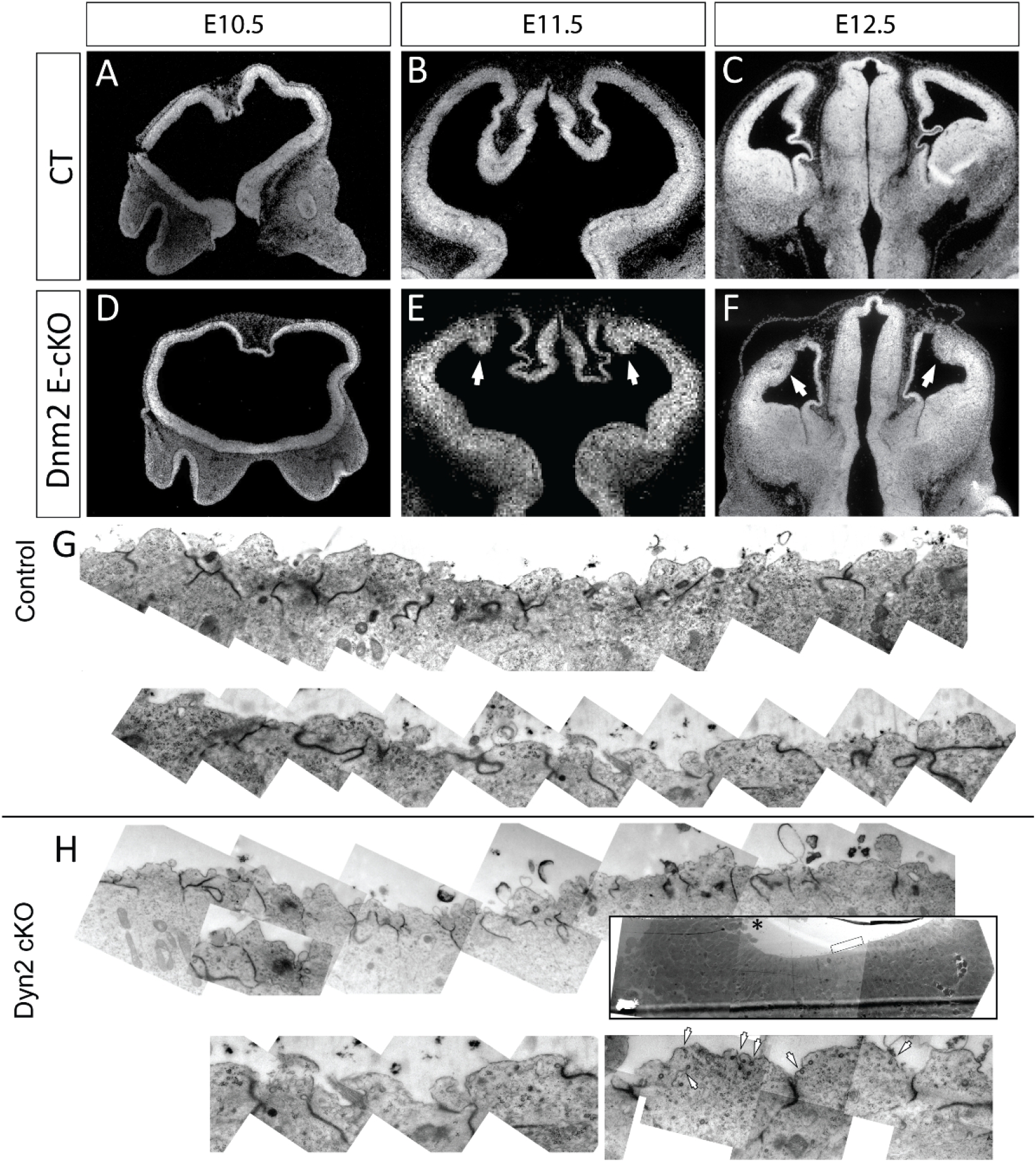
Early disruption of ventricular zone architecture following cortex-specific Dynamin 2 ablation. (A–C) DAPI-stained coronal sections of embryonic forebrain from control (CT) embryos at E10.5 (A), E11.5 (B), and E12.5 (C). Control cortices show normal ventricular zone (VZ) organization, smooth ventricular surfaces, and progressive cortical expansion. (D–F) DAPI-stained coronal sections of embryonic forebrain from Emx1-Cre; Dnm2fl/fl (Dnm2 E-cKO) embryos at E10.5 (D), E11.5 (E), and E12.5 (F). At E10.5, overall cortical morphology appears grossly preserved. By E11.5, focal disruptions of the ventricular surface and abnormal nuclear organization become evident in the dorsal cortex (arrows). By E12.5, these defects expand laterally and deepen, with pronounced ventricular surface irregularities and protrusion of cellular masses into the ventricular lumen (arrows). (G-H) Electron microscopic analysis showing increased number of clathrin coated vesicles and tubular extensions attached to the VZ surface in cKO animals at E11.5.

Immunohistochemistry with cell markers at E10.5 shows a normal population of Sox2+ progenitors in the mutants. At this age, there is heterogeneity in the observations: some mutant animals are indistinguishable from controls, while others start to show an incipient phenotype in the dorsal cortex, where Cre expression corroborates the early gradient of expression described above (**Fig. S3**). The most affected cases show abundant neuronal death (**Fig. S2**) and increased neuronal differentiation in the dorsal cortex, measured by a larger number of intermediate progenitors and early neurons identified by Tbr2, Ngn2 and Tuj1 (**Fig. S2-3**). This increase in the number of Tbr2 and Ngn2 expressing cells sometimes reverses the normal latero-medial gradient observed in controls, as is very clear for Ngn2 (**Fig. S3**).

At E11.5, a depletion of Sox2 + progenitors in the VZ becomes evident (**Fig. 2A-B**), while an abnormally large number of Tbr2+ intermediate progenitors accumulate ectopically around them and even into the ventricle, creating a protrusion in the inner ventricular surface (**Fig. 4A-D**). Abundant pyknotic cells can be detected, and apoptosis is confirmed by cleaved Caspase3 labeling (**Fig. S2**). These data suggest that a defect of Dnm2 affects progenitors behavior by increasing neuronal differentiation. However, to evaluate additional effects of the mutation in differentiating neurons, we bred Nex1-Cre animals with Dnm2 floxed animals. The Nex1 (NeuroD6) promoter starts expression around E11.5 and is restricted to committed neuronal progenitors and neurons in the dorsal cortex (Goebbels, Bormuth et al. 2006). Nex1-Cre Dnm2 cKO animals showed Cre expression in developing neurons at E14 (**Fig. S4**), but no defects or developmental abnormalities could be detected at E14 or later in adult animals (**Fig. S4**). These results confirmed that the primary target of the Dnm2 mutation is the VZ progenitors.

**Figure 4.**
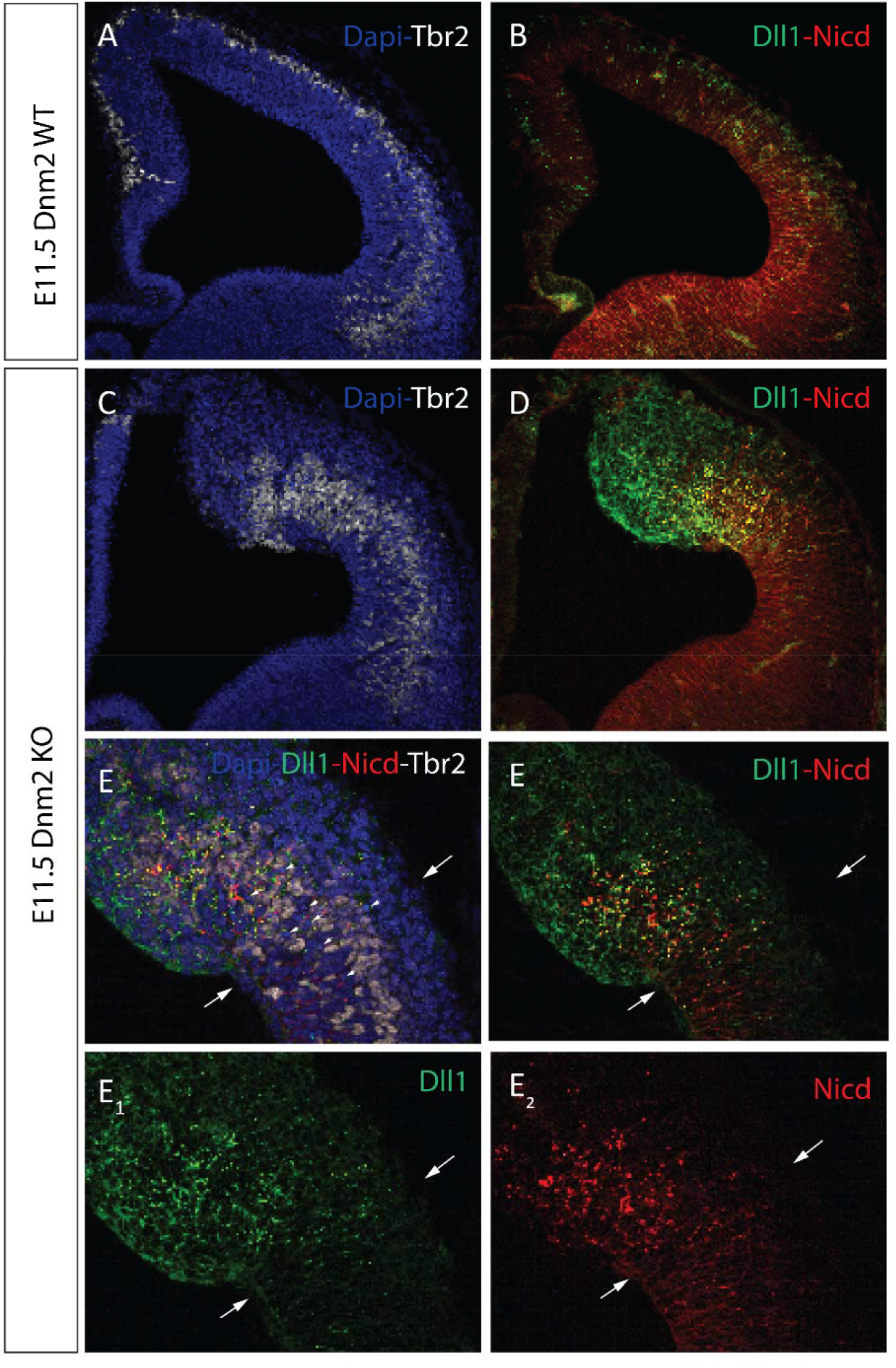
Premature neurogenesis coincides with abnormal accumulation of the Notch ligand Dll1 in the ventricular zone. (A,B) Confocal images of E11.5 control cortex immunostained for Tbr2 and DAPI (A), and Dll1 (green) and NICD (red) (B). Tbr2^+^ intermediate progenitors are largely confined to the subventricular zone, while Dll1 and NICD show diffuse distributions. (C,D) Corresponding images from Dnm2 cKO cortex showing expansion and mislocalization of Tbr2^+^ cells into the ventricular zone and marked accumulation of Dll1 signal. (E) Higher-magnification view of Dnm2 cKO cortex showing ectopic Tbr2^+^ clusters (arrowheads) and regions of intense Dll1 accumulation (arrows). (E_1_, E_2_) Single-channel views of Dll1 and NICD. Dll1 signal is strongly enriched in discrete ventricular regions, whereas NICD distribution is not proportionally increased in nuclei.

Taken together, both the Nestin and the Emx1 driven Dnm2 conditional KOs exhibited a pattern of early differentiation of progenitors. This is supported by a depletion of the VZ progenitors while there are abnormally large numbers of Tbr2 and Tuj1 positive cells in the developing cortex.

Premature depletion of VZ RG could be due to several reasons. A lack of adhesion could cause RG detachment into the cortical plate. This typically leads to the formation of rosette structures (Kadowaki, Nakamura et al. 2007) which we did not observe in this tissue as we had seen in NestinCre; Numb^fl/fl^; Numblike^null/null^ mice (Rasin, Gazula et al. 2007). Further, we noted that the beta-catenin-labeled adherens junctions were largely intact in areas where premature neurogenesis was not observed (**Fig. S5**). This suggests that RG might improperly self-renew, leading to premature depletion because of increased neuronal differentiation of daughter cells.

Thus, loss of Dynamin 2 results in progressive and penetrant defects in cortical morphology across multiple embryonic stages, consistent with early disruption of progenitor homeostasis rather than isolated patterning defects.

### Notch signaling is decreased in Dnm2 cKO mice

Overall, the increase in differentiation of progenitors towards a neuronal fate suggests a Notch phenotype. Therefore, we used antibodies directed against the intracellular domain of the Notch1 receptor (NICD), against the major Notch ligand, Delta-like 1 (Dll1) and against the intracellular domain of Notch—but which will also detect the full length Notch1 receptor. (Notably, this antibody often fails to illuminate the nuclear form of NICD due the instability of this form and tight chromatin association). At E11.5 NICD is widely distributed in the VZ, with cytoplasmic or membrane localization, and exhibiting a marked latero-medial gradient of intensity (**Fig. 4A-B**). Dll1 immunostaining is also widespread, without a clear gradient, and appears mostly as intracellular puncta with several filamentous stains located predominantly in the basal aspect of the cortical primordium (**Fig. 4A-B**). In mutants, there are two clear territories, a lateral aspect where the labeling is similar to that of the controls, and the medial aspect that exhibits the damage (**Fig. 4A-E**). Both Dll1 and NICD appear to accumulate in large and widespread aggregates around the remaining progenitors and differentiating cells in this damaged area. And particularly Dll1 shows a striking accumulation, several fold that of the normal areas (**Fig. 4A-E**). Interestingly, Dll1 and NICD frequently accumulated in overlapping regions, consistent with impaired productive signaling. These results reveal that Dynamin 2 is required to maintain the normal spatial relationship between Dll1 expression, cleaved NICD production, and the balance of RG and progenitors in the embryonic cortex.

To directly assess the level of activation of Notch we evaluated the presence of the active form of NICD (cleaved NICD; heretofore cNICD) which detects the cleavage site epitope around Val1744. (Note that this antibody requires antigen retrieval to detect the unstable cNICD formed by the terminal cleavage event and does not label the vesicular forms seen with the above NICD antibody, which can label all forms of Notch1—cleaved and uncleaved.). cNICD immunostaining demonstrated an uneven staining in the nuclei of VZ progenitors in the cortical primordium (**Fig. 5A-B**). Similar to results with the NICD-targeted antibody, it shows normally a latero-medial gradient of intensity, such that lateral regions show more intense staining than medial ones (**Fig. 5**). However, Dnm2 mutants showed a marked decrease of cNICD in medial regions, corresponding mostly to the areas most affected by the phenotype, but also showing adjacent, more lateral areas with a marked decrease in NICD activation (**Fig. 5B**). Collectively, these data demonstrate that spatiotemporal Dnm2 cKO correlates with decreased cNICD and exuberant neuronal differentiation associated with neurogenic LOF notch phenotypes.

**Figure 5.**
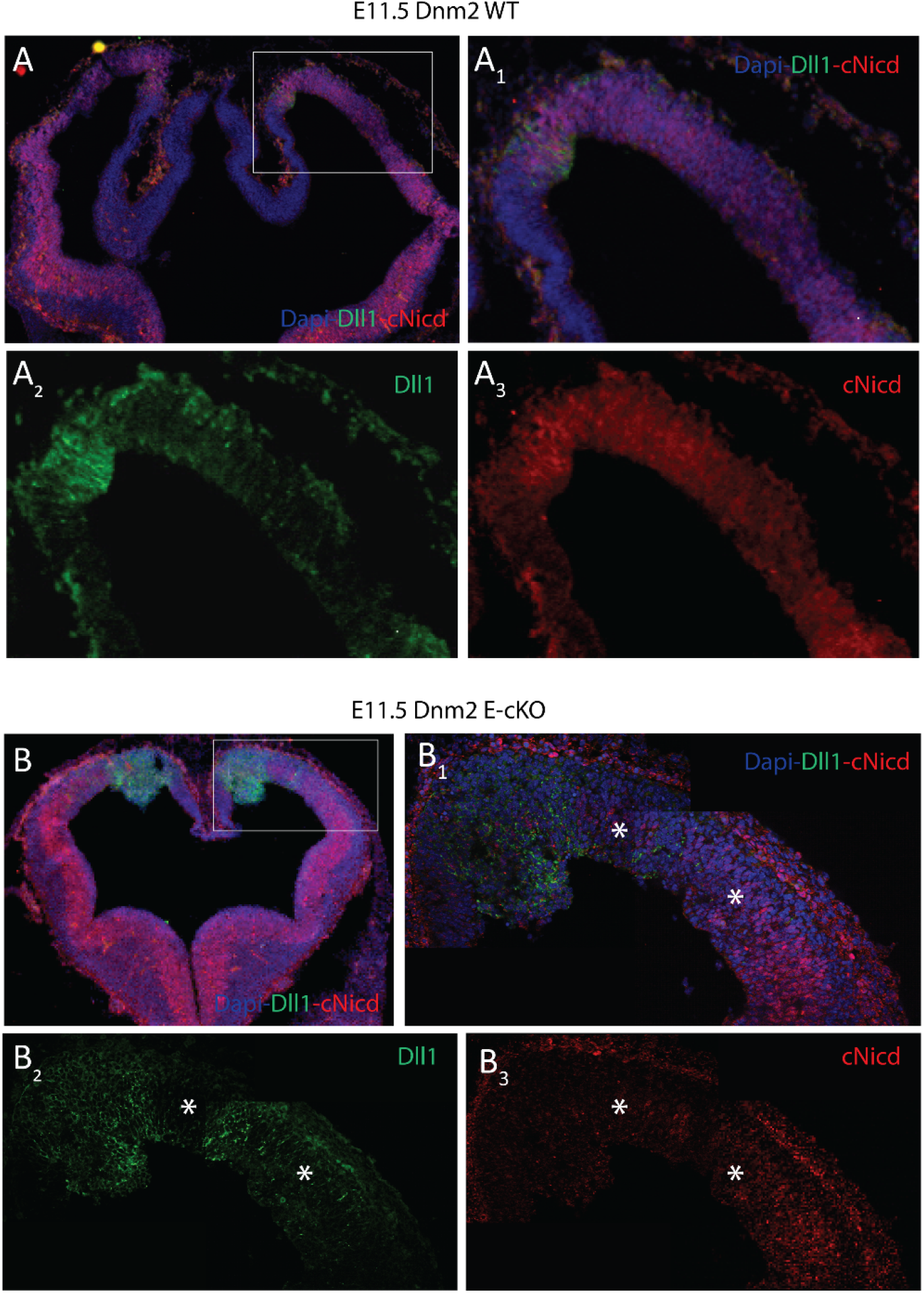
Loss of Dynamin 2 leads to abnormal accumulation of Dll1 and altered NICD distribution in the E11.5 cortex. (A)Low-magnification confocal image of an E11.5 control (Dnm2 WT) coronal cortical section immunostained for Dll1 (green), NICD (red; total Notch intracellular domain), and DAPI (blue). A boxed region indicates the area shown at higher magnification in (A_1_–A_3_). (A_1_) Higher-magnification view of the boxed region in (A), showing the merged DAPI/Dll1/NICD channels in control cortex. (A_2_, A_3_) Single-channel views of Dll1 (A_2_) and NICD (A_3_) corresponding to (A_1_). In control cortex, Dll1 exhibits a punctate intracellular distribution, while NICD is broadly distributed with cytoplasmic and membrane-associated localization throughout the ventricular and subventricular zones. (B) Low-magnification confocal image of an E11.5 Dnm2 E-cKO coronal cortical section immunostained for Dll1, NICD, and DAPI. A boxed region indicates the area shown at higher magnification in (B_1_–B_3_). (B_1_) Higher-magnification merged image of the boxed region in (B). Affected regions of the ventricular zone show striking accumulation of Dll1 signal and altered NICD distribution (asterisks). (B_2_, B_3_) Single-channel views of Dll1 (B_2_) and NICD (B_3_) corresponding to (B_1_). In Dnm2 E-cKO cortex, Dll1 accumulates at markedly elevated levels in discrete ventricular regions, while NICD signal persists but does not show a corresponding increase in nuclear enrichment (asterisks).

Notch signaling contributes to preserve the progenitor fate by inducing expression of target genes like Hes1 and Hes5. We further evaluated the output of Notch signaling by investigating the expression of Hes1. In controls, Hes1 has a similar distribution as cNICD, exhibiting a salt and pepper staining that reflects the lack of expression in non-VZ progenitors, such as Tbr2 positive cells and those downstream in the neuronal fate (**Fig. 6A**). In Dnm2 E-cKOs, a similar pattern can be observed in the lateral, preserved regions of the cortical primordium. However, a notable decrease is detected already at E10.5 in the medial regions, where Dnm2 is starting to be downregulated (**Fig. 6A**). This trend only grows bigger as development proceeds and by E12.5 it is evident that although there is expression of Hes1 in the remaining progenitors, their number, and the intensity is strongly reduced (**Fig. 6B**). In addition, qRT-PCR of Hes5, the other well-characterized target of canonical Notch signaling, indicated a dramatic decrease in normalized mRNA abundance in the mutant CNS. Interestingly, there was a trend toward decreased Notch1 mRNA at the same time but this did not reach significance (**Fig. 6C**). This could be partly explained by the fact that Notch1 is also known to be expressed by differentiating neurons later during their maturation (Sestan, Artavanis-Tsakonas et al. 1999, Breunig, Silbereis et al. 2007). Nevertheless, consistent with disrupted Notch pathway regulation, Dynamin 2 loss results in reduced Hes1 and Hes5 expression, linking altered endocytic dynamics to transcriptional outputs of Notch signaling.

**Figure 6.**
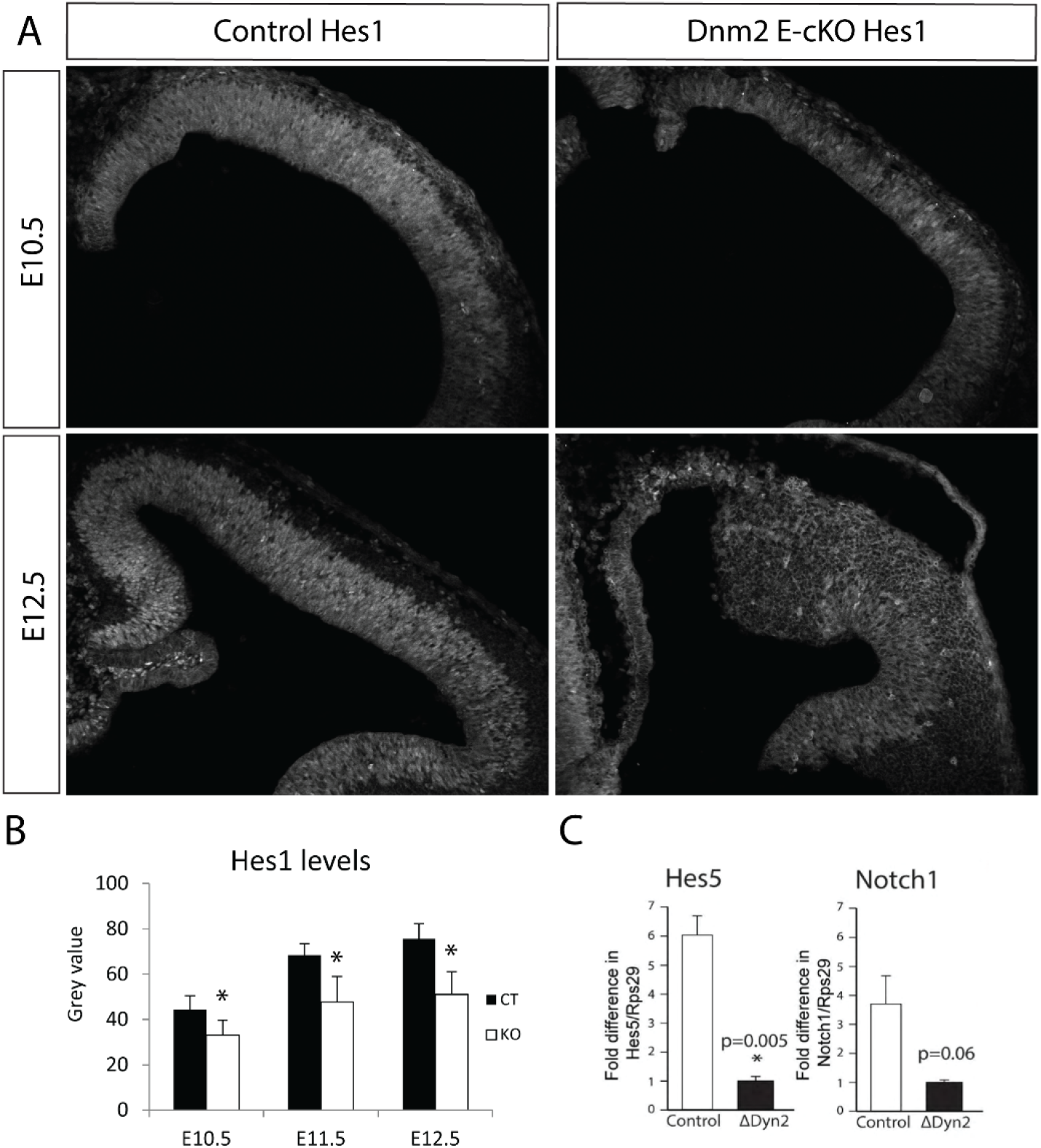
Loss of Dynamin 2 reduces canonical Notch transcriptional output in the embryonic cortex. (A) Confocal images of Hes1 immunostaining in control and Dnm2 cKO cortex at E10.5 and E12.5. Hes1 expression is robust in control ventricular zone and reduced and disorganized in Dnm2 cKO tissue. (B) Quantification of Hes1 immunofluorescence intensity in the ventricular zone across developmental stages. Hes1 levels are significantly reduced in Dnm2 cKO cortex (mean ± SEM; p < 0.05). (C) qRT–PCR analysis of Hes5 and Notch1 mRNA levels in control and ΔDnm2 forebrain. Hes5 expression is significantly reduced, while Notch1 shows a non-significant downward trend.

### Dynamin 2 LOF has opposing effects in Notch signaling and receiving cells in vitro

Reduced Notch activation could reflect three non-mutually exclusive mechanisms. First, reduced Notch signaling activity could be a result of a lack of Dll1 endocytosis and thus a lack of Notch1 receptor dissociation in a non-cell autonomous fashion (Meloty-Kapella, Shergill et al. 2012). Alternatively, previous studies in fly have indicated that Notch internalization is a necessary prerequisite for gamma secretase cleavage (Vaccari, Lu et al. 2008). Thus, the diminished NICD accumulation we observed could be due to a cell autonomous lack of Notch endocytosis and an inability of gamma secretase to cleave the dissociated form of Notch in an intracellular endocytic compartment. Finally, it could result from both of the aforementioned processes (i.e. lack of Notch and Dll1 endocytosis) occurring simultaneously.

To address this question we created a neural stem cell based co-culture system. One population (i.e. Notch “receiving” cells) was nucleofected with full-length Notch1 with a myc tag, a TagBFP blue nuclear reporter protein, a Notch-depended firefly luciferase reporter (CSL-luc), and a Renilla transfection control (**Fig. 7A**). The co-cultured population (Dll1 “sending” cells or controls) was nucleofected with either EGFP (control) or EGFP with Dll1 ligand (“sending,” **Fig. 7A**). Immunostaining demonstrated a high level of expression of the overexpressed proteins and intermixing between the two co-cultured neural stem cell populations (**Fig. 7B**). Cells were allowed to interact for three days prior to lysis and assessment of luciferase activity (**Fig. 7C**). Notably, Dll1 expression in the ligand population created a significant increase in the ratio of firefly luciferase expression when compared with an EGFP-only population (**Fig. 7D**). This increase in luciferase was abrogated by the co-expression of Dynamin 2 K44a-EGFP (heretofore referred to as K44A), a dominant negative form of the protein, in the ligand population; or by expression of a Dnm2 shRNA (**Fig. 7D**). Interestingly, co-expression of K44A in the receptor population (i.e. the Notch1 receptor expressing “receiving” population) caused a significant *increase* in luciferase activity when co-cultured with Dll1 expressing cells (**Fig. 7C**), suggesting that Dnm2-dependent internalization of Notch is not essential for receptor activation. Previous findings have suggested that a form of Notch which lacks the extracellular domain (EMV) but is constitutively cleaved by gamma secretase, requires Dynamin-dependent internalization for activation at the membrane (Gupta-Rossi, Six et al. 2004). However, using this EMV Notch, we failed to find a requirement for Dynamin function in the activation of this truncated Notch receptor in neural stem cells (**Fig. S6**). Specifically, cells transfected with Dynamin K44A and EMV Notch showed a similarly increased level of Notch activity when compared with EGFP and EMV transfected controls (**Fig. S6**).

**Figure 7.**
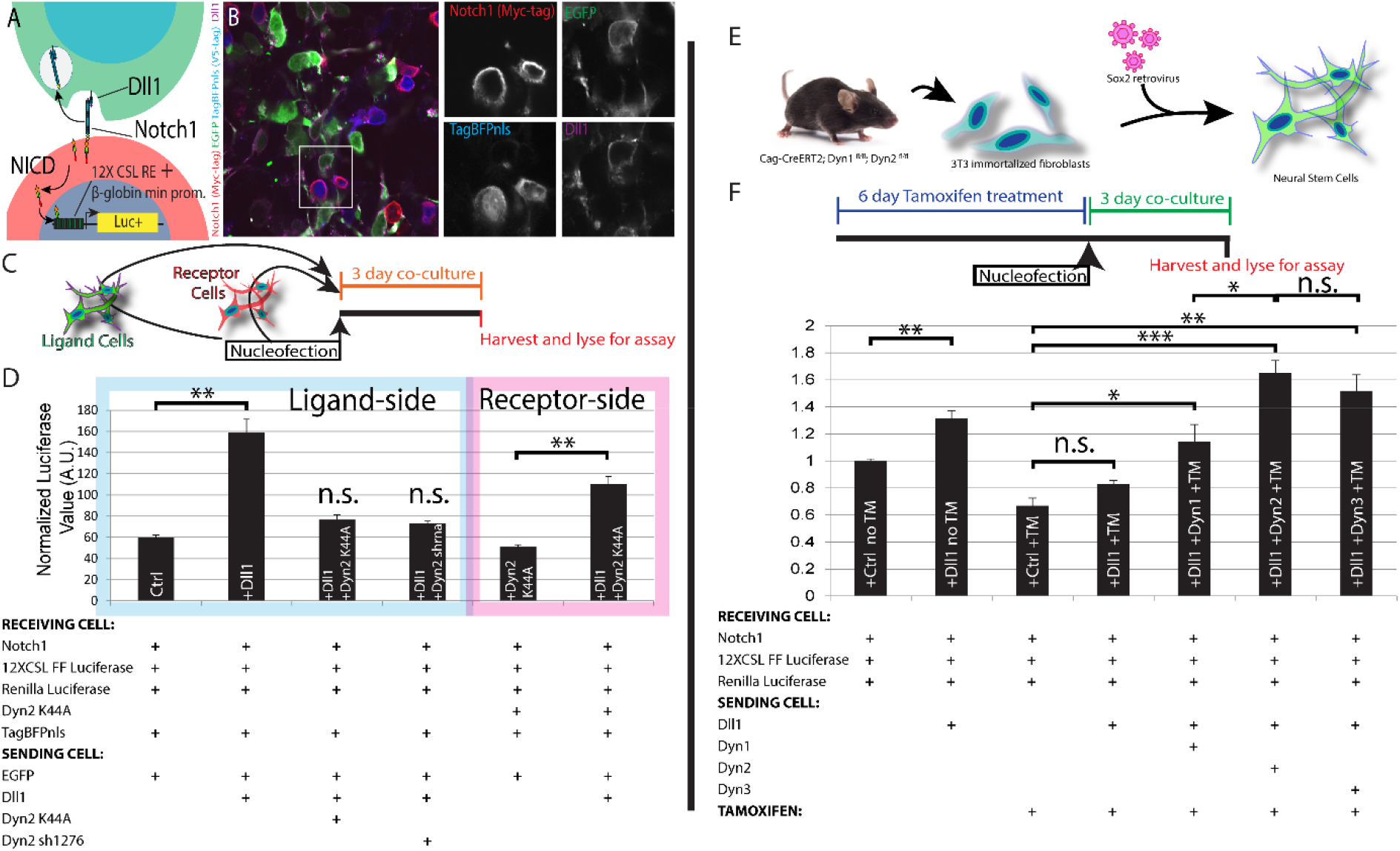
Dynamin 2–dependent endocytosis is required in ligand-presenting cells for efficient Notch activation. (A) Diagram illustrating the Notch signaling assay used throughout this figure. Ligand (Dll1) binding to Notch1 results in release of NICD, which activates a 12× CSL-responsive firefly luciferase reporter. Luciferase activity therefore provides a quantitative readout of Notch signaling strength. (B) Representative images showing expression and localization of Notch1, Dll1, and reporter components in co-cultured cells. (C) Co-culture design distinguishing ligand (“sending”) and receptor (“receiving”) cell populations. (D) Quantification of Notch-dependent luciferase activity. Inhibition of Dnm2 in ligand-presenting cells (K44A or shRNA) reduces Dll1-mediated activation, whereas inhibition in receptor-expressing cells increases signaling. (E) Generation of inducible Dnm2-deficient neural stem cells by Sox2-mediated reprogramming. (F) Rescue assay showing restoration of Notch signaling by re-expression of dynamin isoforms following Dnm2 deletion. (G) Quantification of rescue experiments demonstrating partial functional redundancy among Dynamin 1, 2, and 3. Statistical significance is indicated; n.s., not significant.

To interrogate the functional equivalence of the Dynamin isoforms, we reprogrammed fibroblasts into neural stem cells from a tamoxifen-inducible CAG-ERT2; Dnm2^fl/fl^ mouse line using Sox2 (**Fig. 7E**) (Ring, Tong et al. 2012). Adapting our co-culture system, we noted that loss of Dynamin 2 significantly abrogated Notch reporter activity in Notch1-expressing cells when we inducibly ablated Dyn 2 in Dll1-expressing cells. As expected, Dynamin 2 overexpression was able to rescue the loss of Dynamin 2 from the floxed locus **(Fig. 7F**). Also notable was the ability of Dynamin 1 and 3 to rescue Dynamin 2 ablation to varying but significant degrees (**Fig. 7F**). These coculture and reporter assays demonstrate that Dynamin activity is differentially utilized in ligand-sending and signal-receiving cells to support physiological Notch pathway activation.

### Interrogation of Dynamin 2 loss of function in vivo results in differential effects on Notch signaling in receptor versus ligand cells

Taken together, our evidence suggests the necessity of intact Dynamin function in ligand presenting cells and fails to support a similar role in the receptor cells. However, the argument has been made that overexpressing Notch presents a non-physiological situation that is not Dynamin-dependent or these results could represent a cell culture or overexpression artifact (Vaccari, Lu et al. 2008, Sorensen and Conner 2010). Thus, we sought to assess the role of Dynamin in Notch-expressing cells in vivo in their natural physiological context. We employed electroporation of the postnatal ventricular zone as a means to interrogate Notch-mediated RG maintenance in a more facile manner (Breunig, Levy et al. 2015, Kim, Rincon Fernandez Pacheco et al. 2019). Specifically, postnatal electroporation allows for comparisons of multiple control and experimental groups within single litters of animals(Kim, Rincon Fernandez Pacheco et al. 2019). Secondly, the protracted cell cycle length allows for more precise manipulation of stem and progenitor cells in the context of Cre recombination(Kim, Rincon Fernandez Pacheco et al. 2019). For example, we have observed that it can take up to six days to deplete Dnm2 following Cre expression (Ferguson, Raimondi et al. 2009). This is comparable to the entire period of cortical neurogenesis whereas the postnatal VZ is a continuous site of Notch-mediated neuro- and gliogenesis, allowing for protracted examination of perturbations of Notch signaling (Ables, Breunig et al. 2011). To ascertain the relative Notch activity of neural stem and progenitors in the region, we created a dual reporter assay utilizing well-characterized Dll1 and Hes5 promoter/enhancer elements (Beckers, Caron et al. 2000, Ohtsuka, Imayoshi et al. 2006) driving destabilized, membrane-tagged mOrange2 and destabilized EGFP, respectively (**Fig. 8A**). (A blue fluorescent nuclear protein, TagBfp2-V5-nls, driven by the constitutive chick beta actin promoter served as an electroporation control [**Fig. 8A**]). When compared with controls, which displayed a mix of Hes5-EGFP+ and Dll1-mOrange2+ cells (**Fig. 8B-F**), dominant negative abrogation of Dnm2 function caused consistent but insignificant increase in Hes5-EGFP+ cells, and a significant decrease in the percentage of Dll1-mOrange2+ cells (**Fig. 8C, F**). A decrease in the percentage of Dll1-mOrange2+ cells was also observed in animals expressing an Rbpj-VP16 fusion protein (**Fig. 8D, F**), which has previously been shown to activate canonical Notch signaling (Mizutani, Yoon et al. 2007).

**Figure 8.**
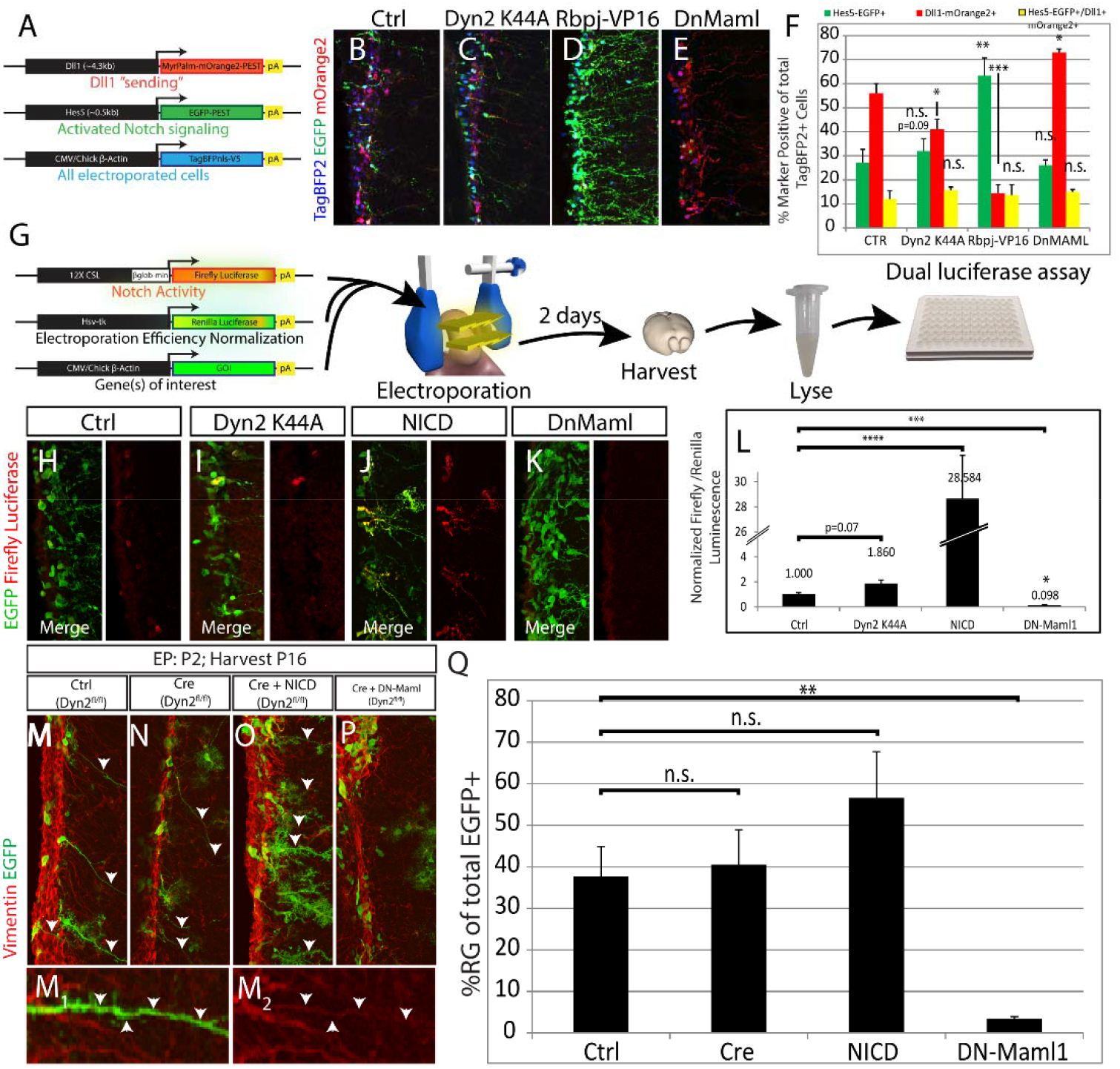
Mosaic Dynamin 2 loss increases Notch activity without depleting radial glial stem cells. (A) Schematic of reporter constructs used for in vivo electroporation. A Dll1 promoter–driven mOrange2 reporter labels ligand-expressing cells, while a Hes5 promoter–driven EGFP reporter marks cells with active Notch signaling. TagBFP2-NLS labels electroporated cells. (B–E) Representative confocal images of the postnatal ventricular zone following electroporation with control (B), Dnm2 K44A (C), Rbpj-VP16 (D), or dominant-negative MAML (DnMaml) (E), showing Hes5-EGFP and Dll1-mOrange2 reporter activity. (F) Quantification of reporter activity. Dnm2 inhibition (K44A) increases Hes5-EGFP^+^ cells without a corresponding increase in Dll1-mOrange2^+^ cells, consistent with enhanced cell-autonomous Notch signaling. (G) Schematic of in vivo dual-luciferase assay for quantitative measurement of Notch signaling. (H–K) Representative images of luciferase reporter expression following electroporation with control, Dnm2 K44A, NICD, or DnMaml constructs. (L) Quantification of normalized firefly/Renilla luciferase activity. Dnm2 K44A increases Notch signaling, whereas NICD activates and DnMaml suppresses reporter activity. (M–P) Confocal images of EGFP^+^ electroporated cells in the ventricular zone stained for Vimentin following control (M), Cre-mediated Dnm2 deletion (N), Cre + NICD (O), or Cre + DnMaml (P) expression. Arrowheads indicate radial glial cells with intact ventricular contact. (M_1_, M_2_) Higher-magnification views showing preserved radial glial morphology following Dnm2 loss. (Q) Quantification of radial glial abundance. Cre-mediated Dnm2 deletion does not significantly reduce radial glial cells, whereas modulation of Notch signaling alters radial glial maintenance (mean ± SEM; n.s., not significant).

However, Rbpj-VP16 also significantly increased the percentage of Hes-EGFP+ cells (**Fig. 8D, F**). Expression of a dominant negative mastermind, which has been shown to attenuate Notch signaling by interfering with formation of the ternary complex of Notch, Rbjk, and Maml (Maillard, Weng et al. 2004), yielded a dramatic increase in Dll1-mOrange2+ cells but did not change Hes5-reporter expression percentages though intensity was notably diminished (**Fig. 8E-F**). To quantitatively measure these changes, we adapted the dual luciferase assay for in vivo use. Specifically, CSL firefly luciferase was electroporated into Notch-dependent postnatal neural stem cells along with a Renilla-expressing plasmid to normalize transfection levels (**Fig. 8G**). Littermate animals were separated into four groups to ascertain the role of Dnm2 blockage on Notch signaling. Notably, overexpression of the Dy2 K44a mutation significantly increased Notch activity in this population of cells in vivo compared with a similar membrane associated EGFP electroporation, in keeping with the previous observations in receptor expressing cells in vitro (**Fig. 7D**) and in the Hes5-EGFP population in vivo (**Fig. 8C and F**). Importantly, this system is able to robustly and reliably readout changes in Notch signaling levels mediated by classically described gain-(e.g. NICD) and loss-of-function (DnMAML) transgenes in a quantitative (**Fig. 8L**) manner. Dominant negative proteins can cause artifactual effects due to overexpression and their ability to sequester binding proteins. Therefore, we electroporated Cre recombinase into homozygous Dnm2 floxed littermates to ablate Dnm2 protein in a cell autonomous manner. RG cell numbers were used as a surrogate for Notch signaling activity given the well-characterized role for the Notch signaling in this process. Notably, there was little difference observed in the percentage of RG between the Ctrl and Cre groups (**Fig. 8M,N, and Q**). (RG were defined by morphology—specifically having a single, long, basal, radial process with a cell body in the VZ—and the expression of the intermediate filament Vimentin in the radial fiber [**Fig. 8M**_1-2_]). In keeping with the ability of Notch to regulate RG cell numbers, NICD and Dn-Maml1 caused opposite effects on the relative percentages of RG with NICD causing consistent but insignificant increases (**Fig. 8O, Q**) while Dn-Maml1 prematurely depleted this stem cell population (**Fig. 8 P-Q**). Taken together, Dynamin 2 loss-of-function appears to *increase* Notch activity cell-autonomously both in vitro and in vivo in receptor-expressing cells. However, this phenotype is only observable in assays which can perceive cell autonomous signaling due to the precise control attainable in vitro or when signaling is manipulated in a mosaic manner in vivo (e.g. electroporation). When the loss of Dnm2 is tissue-wide, the neurogenic loss-of-Notch function predominates due to a lack of ligand activity. Taken together, our findings support a model in which Dynamin2-mediated endocytosis coordinates ligand–receptor trafficking to spatially and quantitatively regulate Notch signaling during cortical development.

## DISCUSSION

We have shown that Dnm2 is essential for the proper cleavage and activation of the Notch receptor in a non-cell autonomous manner by promoting ligand endocytosis. While endocytosis has been associated with Notch signaling, the details (ligand vs. receptor side function) and mammalian Dynamin isoform responsible for the signaling activity have remained unclear. The results here identify Dnm2 as the predominant physiological dynamin supporting Notch signaling in embryonic neural stem and progenitor cells. It will be interesting to learn if this is the case in other tissues—especially given that Dynamin 1 and Dynamin 3 appear to be expressed predominantly in the postnatal brain. (However, embryonic atlases do illuminate some non-CNS populations that are exceptions to this pattern such as mesodermal populations [Supp. Fig. 1A-H]). In addition, given this expression pattern of these latter two isoforms, the question arises as to whether Dnm2 is solely essential for the Notch signaling that occurs at this point in both precursors and neurons (Breunig, Silbereis et al. 2007, Ables, Breunig et al. 2011, Alberi, Liu et al. 2011).

### Role of Dnm2 in Cell Autonomous Notch Cleavage

Previously, the role of Dynamin in the physiological cleavage of Notch has been controversial (Struhl and Adachi 2000, Gupta-Rossi, Six et al. 2004, Vaccari, Lu et al. 2008, Sorensen and Conner 2010). The primary argument has been that many of the negative findings which failed to find a role for Dynamin in Notch activation were performed in vitro in the context of Notch overexpression (Vaccari, Lu et al. 2008). We have explored the role of Dynamin in the context of physiological Notch expression as well as Notch overexpression. Both cases provide evidence that Dynamin LOF increases Notch activity in endogenous neural stem cells in vivo and in vitro. Taken together, this indicates that γ-secretase cleavage can occur at the membrane. Indeed, while γ-secretase cleavage is favored at the higher PH’s which are observed in these compartments—though there may be other properties of these endosomes alter cleavage sites and alter NICD stability (Pasternak, Bagshaw et al. 2003, Kanwar and Fortini 2008, Tagami, Okochi et al. 2008). Elegant studies in Drosophila indicate that proper sorting of receptor and ligands intracellularly is exquisitely complex and likely constitutes another level of regulation of this relatively “simple” pathway (Bray 2006, Fortini and Bilder 2009). However, the finding of alternate cleavage sites with functional implications on NICD stability (Tagami, Okochi et al. 2008) adds another level of complexity with direct relevance to the clinical Notch therapies and associated Notch mutations found in many cancers.

Also, we have described the protein localization of Notch1, and Dll1 in normal embryonic brain and in our Dnm2 mutants which lack a specific form of clathrin-mediated endocytosis. These results suggest a stereotypical, segregated pattern of expression that rapidly changes upon any alteration from steady-state conditions. The immunostaining pattern that we observed is largely in agreement with previous FACS, in situ, and microarray data from several groups, and suggests that progenitor types can be largely defined by expression of Notch signaling components (i.e. precursors express Notch1 receptors and target genes while committed progenitors express ligands and proneural genes which antagonize Notch signaling) (Mizutani, Yoon et al. 2007, Kawaguchi, Ikawa et al. 2008, Nelson, Hodge et al. 2013).

### Pan-CNS Dnm2 Ablation Mimics Notch LOF

The severity of our mutant is profound, leading to late prenatal or early postnatal lethality, and is comparable to other Notch mutants such as nervous system specific Dll1 and Mib1 conditional knockout mice (Kawaguchi, Yoshimatsu et al. 2008, Yoon, Koo et al. 2008). Notably, we observe severe hemorrhaging which is also observed in a CNS-specific Nestin-Cre; Dll1^fl/fl^ mutant (Kawaguchi, Yoshimatsu et al. 2008). Also, at the timeline of progression of the neurogenic phenotype is largely in agreement. However, a few mice appeared to present a more severe phenotype at earlier stages than the Dll1 cKO. This may be due to the fact that Dll1 expression is restricted to INPs and thus recombination in this cell type may take more time. Alternatively, the contribution of other ligands (e.g. Jag1) may attenuate the Dll1 phenotype.

The Dnm2 cKO mice provide a means to block all Notch signaling without altering the repressor function of RBPj. We cannot completely rule out Notch-independent effects of Dnm2 ablation as this protein likely has pleiotropic roles in many signaling pathways. For example, we have noted a rather dramatic cell death phenotype, which has not been described in other Notch mutants nor does it seem to be attributable to the later hemorrhaging. However, as is often seen in Drosophila, the Notch phenotype predominates due to its exquisite dosage sensitivity (Jaekel and Klein 2006, Andersson, Sandberg et al. 2011, Guruharsha, Kankel et al. 2012). It should be noted that chemical inhibitors of Dnm2 have been used to temporarily block protein activity. However, these have been demonstrated to have significant non-specific effects (Park, Shen et al. 2013).

In conclusion, despite the presence of two other mammalian isoforms of Dynamin, we have identified Dnm2 as essential for proper physiological endocytosis of Notch ligands and their non-cell autonomous activation of the Notch receptor. However, we find that loss of Dnm2 in receptor expressing cells functions to increase Notch activity, arguing against the necessity of receptor internalization for Notch activity during physiological Notch cleavage in the development of the mammalian central nervous system. Taken together, Dnm2 is a molecule essential for proper brain development by mediating asymmetric division during precursor cell divisions. These findings establish endocytic asymmetry as a fundamental mechanism for directional Notch signaling, providing a framework for understanding how identical signaling components can produce divergent outcomes depending on cellular context.

## EXPERIMENTAL PROCEDURES

### Mice

Generation of a floxed allele for Dnm2 has been previously described (Ferguson, Raimondi et al. 2009). Time-pregnant litters were collected by cesarian section at the times indicated and tissue was handled as described for each method below.

### Immunohistochemistry

Whole embryos or embryo heads were collected in 4% PFA and fixed for 4-6 hours. Tissue was then transferred to 10% sucrose until it sunk and then was put into 30% sucrose. After sinking, the tissue was snap frozen in OCT and sectioned in a cryostat onto SuperFrost Plus glass slides. For standard immunohistochemistry. Slides were washed 3 times in PBS, then blocked with 3% normal donkey serum in PBS plus triton. Primary antibodies were added to this same blocking solution and the sections were incubated overnight at 4°C. The following day, the sections were washed again three times in PBS. Donkey Secondary antibodies (Jackson Immunoresearch, Dylight 488, Dylight 549, Dylight 649, Biotin or AMCA-conjugated, species-specific, maximally cross-absorbed) were incubated for one hour at room temperature at 1:1000. Sections were then washed three times in PBS and counterstained with DAPI (1:3000). Cleaved NICD required antigen retrieval and tyramide signal amplification. Briefly, after washes and prior to blocking, tissue was incubated in 1X Dako Target Retrieval Solution at 95°C for 20 minutes. It was allowed to cool to room temperature in this same solution. Then, tissue was washed, blocked, and incubated in primary antibody as per normal immunohistochemistry. After overnight incubation, sections were washed and incubated with Donkey anti-rabbit Biotin-conjugated secondary antibody. After washing, tissue was incubated with fluorophore-conjugated strepavidin. Then, tissue was washed in PBS and counterstained.

### qRT-PCR

Cerebral hemispheres were collected into Trizol and immediately homogenized. This solution was kept at -80° C. RNA was extracted according to the manufacturer’s instructions. Genomic DNA was digested for 1 hr at 37° C using Turbo DNAFree. This DNAse was removed according to manufacturer’s instructions and RNA was transferred to a fresh tube and stored at -80° C. cDNA was synthesized using 10 ng RNA, random pentadecamers (IDT) and the Superscript III kit. RNA was then digested using RNase H. 10 ng cDNA was used as a template for qRT-PCR with Taqman probes in an Applied Biosystems 7900HT Real-time PCR system.

### Immunoblotting from embryonic brains

Brain tissue was collected from E18 embryos and frozen by immersion in liquid N_2_. After genotyping, brain samples were thawed in 25 mM Tris + 150 mM NaCL + 1mM EDTA +1% Triton X-100 + protease inhibitor Cocktail (Roche), sequentially homogenized with a Potter Elvehjem homogenizer followed by a Qiashredder column (Qiagen). Proteins were resolved on 8% SDS-PAGE gels, transferred to nitrocellulose and probed with previously described and validated antibodies (Ferguson, Brasnjo et al. 2007).

### Neural Stem Cell Culture

Neural stem cells were harvested as previously described(Breunig, Silbereis et al. 2007). Cells were cultured continuously as monolayers in 20 ng/ml FGF-2, 20 ng/ml EGF, and 5 ng/ml heparin in Neurobasal medium plus B27 in the presence of antibiotic-antimycotic. For immunostaining, cells were transferred mechanically into 24 well plates containing growth-promoting coverslips (FISHER). After a brief wash in PBS with calcium and magnesium, cells were fixed with 4% PFA for 15 minutes.

### Luciferase Assay

Following nucleofection according to manufacturer’s directions (10 ug of DNA per 5 × 10^6 cells), cells were allowed to grow in flasks for 2 days. Cells were lysed according to the manufacturer’s instructions (Promega Dual-Glo) and results were read with a Envision 2104 Plate plate reader. Renilla luciferase served as a normalization control. Luminescence was measured using a plate reader and expressed as normalized Firefly/Renilla ratios.

### Tamoxifen Administration

For inducible recombination experiments, tamoxifen (Sigma) was dissolved in corn oil (20 mg/mL) and administered at **100 mg/kg/day** for **5–6 consecutive days**, followed by a **3-day chase period** prior to tissue collection or coculture assays (Figures 7–8).

### Imaging and Quantification

Images were acquired using **confocal microscopy** (Zeiss LSM series or Nikon A1R) or Zeiss Apotome structured illumination. Identical acquisition settings were used for control and mutant comparisons.

Cell counts were performed on matched cortical regions across genotypes and developmental stages.

### Postnatal Electroporation

Postnatal electroporation was conducted between P1-P2, targeting the ventricular zone of the lateral ventricles (Breunig, Levy et al. 2015). Plasmids were injected at 1–2 µg/µL followed by square-wave electrical pulses.

### Transcriptomic Data Sources

Publicly available single-cell and spatial transcriptomic datasets were downloaded from the following sources: Mouse embryonic and early postnatal single-cell RNA-seq datasets spanning E8.5–P0 were obtained from publicly available resources generated by the **Shendure laboratory (https://omg.gs.washington.edu/)**. Single-cell RNA sequencing datasets used for lineage and dynamin co-expression analysis were obtained from **Parse Biosciences–generated E18 Mouse Brain datasets. Spatial transcriptomics datasets 10x Genomics Visium HD** mouse brain datasets (E15.5 cortex) were downloaded directly from the **10x Genomics public data repository**.

### Single-Cell RNA-Seq Processing and Analysis

Single-cell RNA sequencing data were processed using **Seurat (v4+)** in R and **Scanpy** in Python. Raw count matrices were filtered to remove low-quality cells and genes based on standard quality control metrics, including total UMI counts, number of detected genes, and mitochondrial transcript percentage.

Data were normalized using log-normalization, scaled, and subjected to dimensionality reduction using principal component analysis (PCA). Neighborhood graphs were constructed using the top principal components, and clustering was performed using the **Leiden algorithm**. Uniform Manifold Approximation and Projection (**UMAP**) was used for visualization.

### Spatial Transcriptomics (Visium HD) Analysis

Visium HD spatial transcriptomics data were processed using **Seurat** with spatial extensions. Gene expression matrices were normalized and mapped to tissue coordinates provided by 10x Genomics.

Spatial feature plots were generated to visualize regional expression of **Sox2, Rbfox3, Dnm1, Dnm2**, and **Dnm3** across the developing cortex. Anatomical regions were annotated based on spatial expression patterns and reference atlases.

### Data Visualization

All UMAP, feature, and spatial plots were generated using **Seurat, Scanpy**, and custom visualization scripts in R and Python. Identical color scales were used for direct comparison of gene expression across datasets where applicable.

## SUPPLEMENTAL DATA

The Supplemental Data for this article can be found online at [insert link]

## ACKNOWLEDGEMENTS

We thank P. Camilli and S.M. Ferguson for the gift of the Dynamin 2 floxed mice, tissues, and data. This work was supported by grants from the National Institutes of Health, including NINDS (R01NS131782; Contact PI J.J.B.) and NCI (R33CA278564). Additional funding was provided by the U.S. Department of Defense (NF220040), CureSearch for Children’s Cancer (Acceleration Initiative Award), and the American Cancer Society Research Scholar Award (all to J.J.B). Support was also received from Cedars-Sinai Medical Center, including funds from the Samuel Oschin Comprehensive Cancer Institute and the Board of Governors Regenerative Medicine Institute.

We acknowledge core facilities and shared resources at Cedars-Sinai Medical Center for imaging, histology, and animal care support. We thank current and former members of the Breunig laboratory for technical assistance, experimental discussions, and critical feedback.

The content is solely the responsibility of the authors and does not necessarily represent the official views of the National Institutes of Health, the Department of Defense, or other funding agencies.

## SUPPLEMENTARY FIGURES

**Figure S1.**
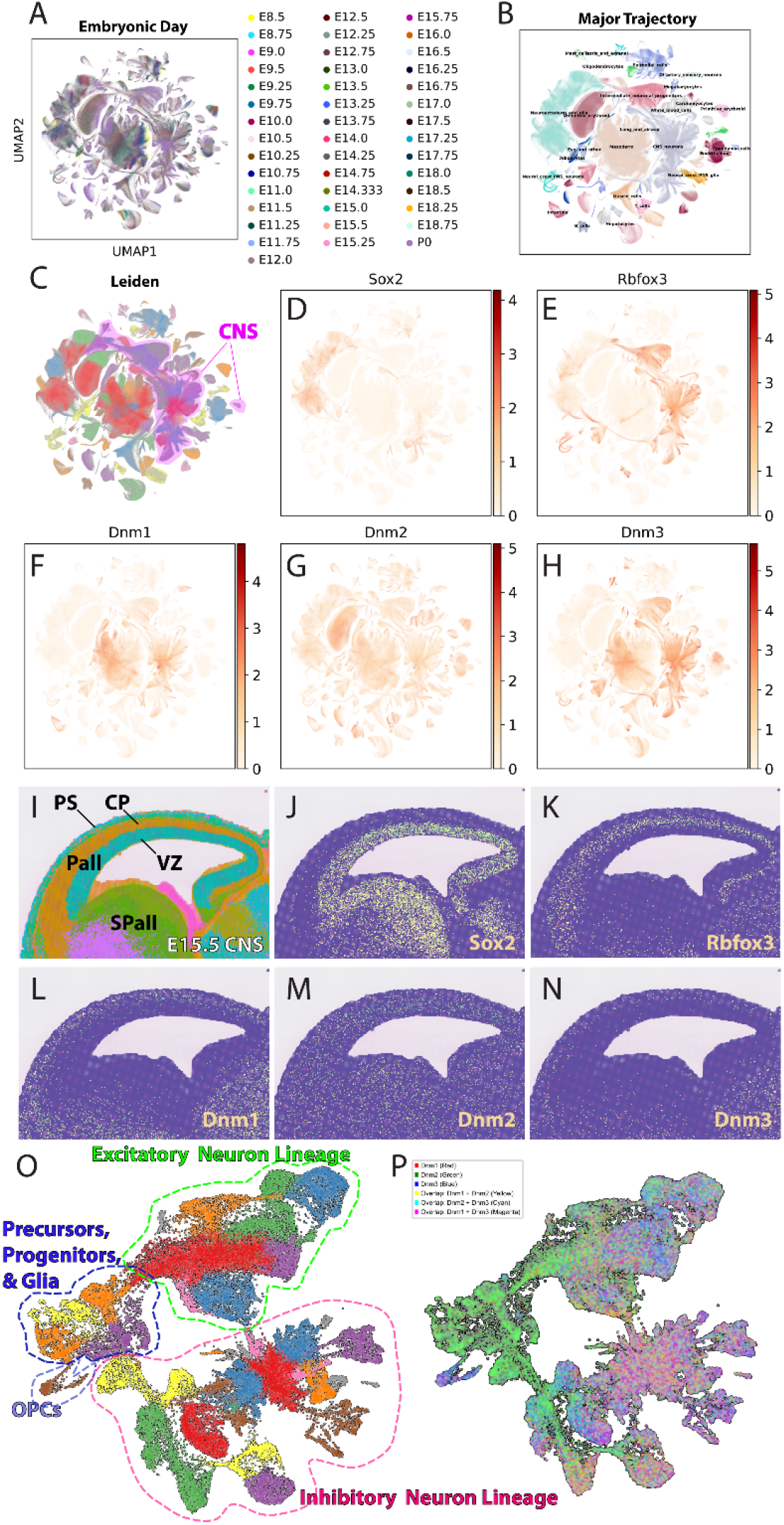
Developmental and lineage-specific expression of Dynamin genes in the embryonic CNS. **(A)** UMAP projection of single-cell RNA sequencing data spanning embryonic day (E)8.5 through postnatal day 0 (P0), colored by embryonic age, illustrating continuous developmental progression across cell states. **(B)** Major developmental trajectories inferred from the integrated dataset, with annotated cell classes including neural progenitors, intermediate progenitors, excitatory and inhibitory neurons, glial lineages, and non-neural cell types. **(C)** Leiden clustering of all cells, highlighting central nervous system (CNS) populations used for subsequent analyses. **(D–E)** Feature plots showing expression of **Sox2** (neural stem and progenitor marker) and **Rbfox3 (NeuN)** (postmitotic neuronal marker), confirming expected distribution across progenitor and neuronal populations. **(F–H)** Feature plots depicting expression of **Dnm1, Dnm2**, and **Dnm3**, respectively, across all cell populations. All three Dynamin genes are broadly expressed, with overlapping but distinct enrichment patterns across neural lineages. **(I)** Visium HD Spatial mapping of annotated progenitor domains onto an E15.5 CNS reference, indicating ventricular zone (VZ), subventricular zone (SVZ), cortical plate (CP), pallium (Pall), and subpallium (SPall). **(J–K)** Visium HD –based expression of **Sox2** (J) and **Rbfox3** (K) at E15.5, demonstrating progenitor-restricted and neuron-restricted expression, respectively. **(L–N)** Visium HD showing expression of **Dnm1, Dnm2**, and **Dnm3** at E15.5, revealing widespread expression throughout the developing cortex with detectable signal in both progenitor and neuronal compartments. **(O)** UMAP highlighting major neural lineages, including progenitors and glia (blue dashed outline), excitatory neuron lineage (green dashed outline), inhibitory neuron lineage (pink dashed outline), and oligodendrocyte precursor cells (OPCs). **(P)** Overlay of **Dnm1** (red), **Dnm2** (green), and **Dnm3** (blue) expression across the UMAP, illustrating extensive co-expression with lineage-specific differences in relative enrichment. Note the lack of Dnm1/3 in the primary precursor/progenitor/glia compartment, the enrichment of Dnm1/Dnm3 in OPCs and the increase in these mRNAs in differentiated neurons.

**Figure S2.**
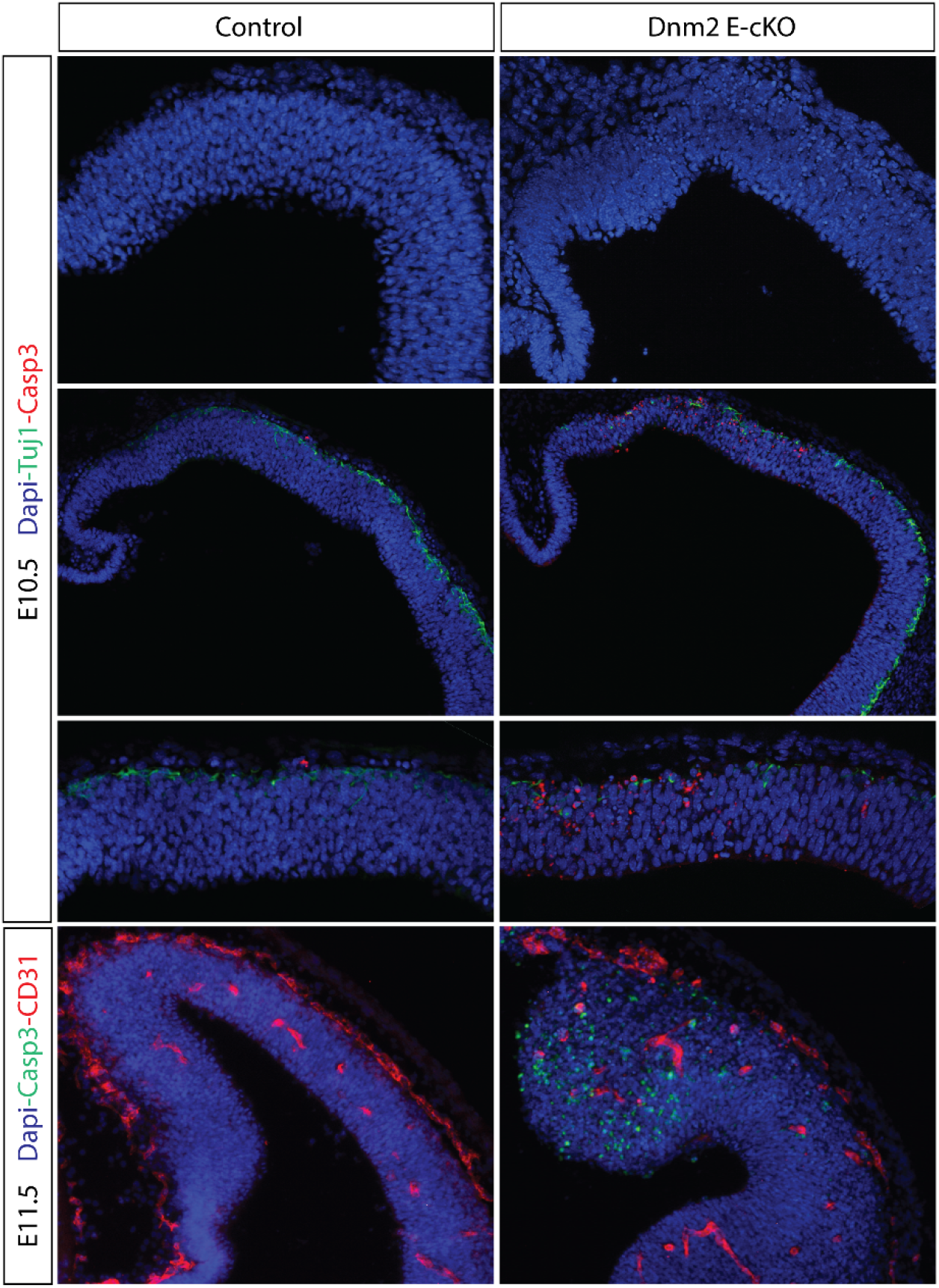
Increased apoptosis and vascular abnormalities following early loss of Dynamin 2 Immunofluorescence analysis of control and Dnm2 E-cKO embryonic cortex at E10.5 and E11.5. At **E10.5** (top three rows), sections were stained with **DAPI** (blue) to label nuclei, **Tuj1** (green) to mark early postmitotic neurons, and **cleaved Caspase-3 (Casp3)** (red) to detect apoptotic cells. Control cortices show sparse Casp3-positive cells, whereas **Dnm2 E-cKO** cortices exhibit increased numbers of Casp3-positive cells distributed throughout the neuroepithelium, particularly within ventricular and subventricular regions. At **E11.5** (bottom row), sections were stained with **DAPI** (blue), **cleaved Caspase-3 (Casp3)** (green), and **CD31** (red) to visualize apoptotic cells and vascular endothelium, respectively. Compared to controls, **Dnm2 E-cKO** cortices display elevated apoptosis accompanied by disrupted cortical morphology and altered vascular patterning. Images are representative of multiple embryos per genotype. Identical imaging and processing parameters were used for control and mutant samples.

**Figure S3.**
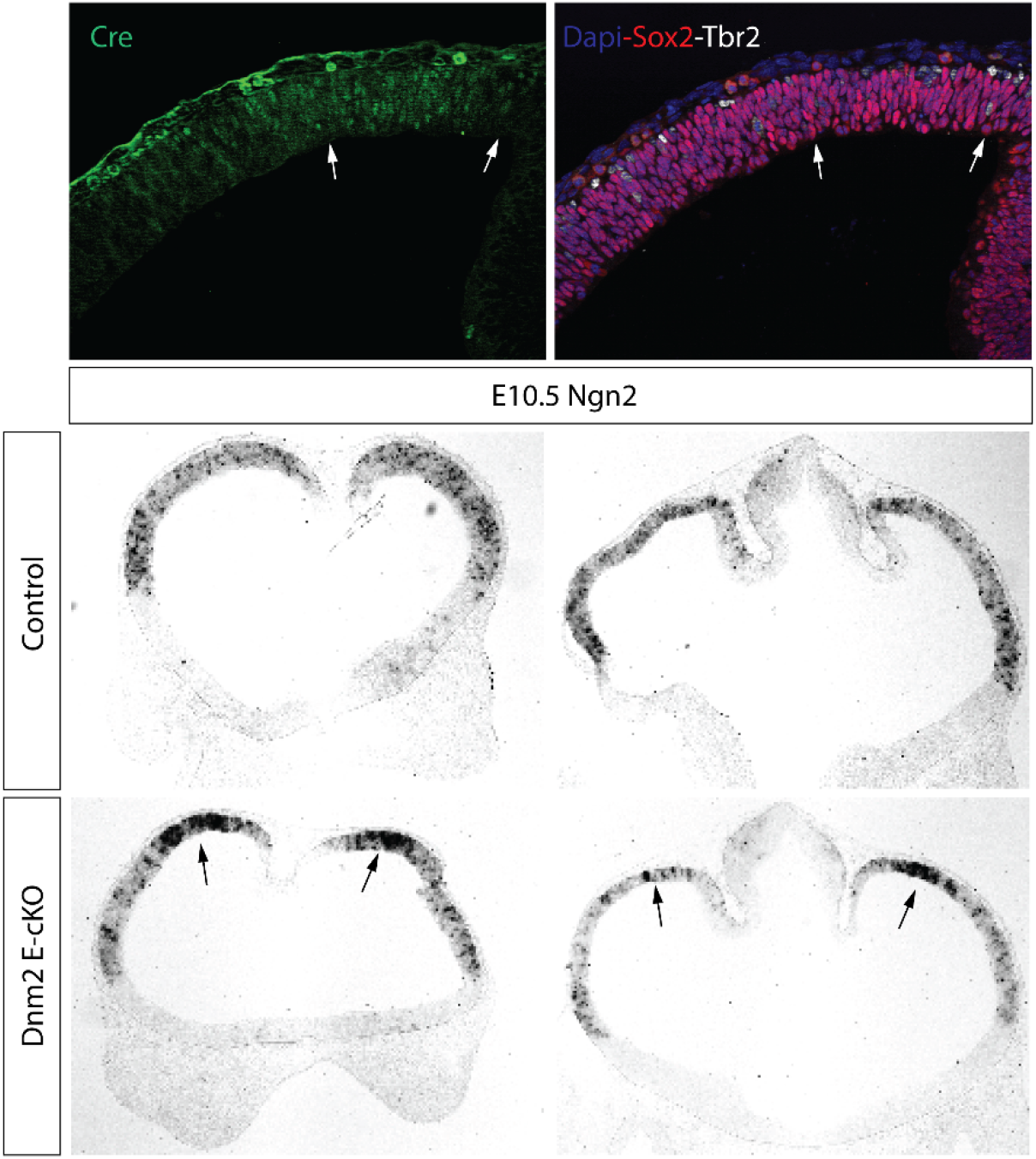
Early neuronal differentiation markers are altered following Dnm2 deletion. **Top row**: Immunofluorescence analysis at **E10.5** showing **Cre** expression (green) in the dorsal telencephalon, confirming efficient recombination in the neuroepithelium. Adjacent sections are stained with **DAPI** (blue), **Sox2** (red), and **Tbr2** (white) to visualize radial glial progenitors and intermediate progenitors, respectively. Arrows indicate regions of altered progenitor organization in Cre-expressing domains. **Bottom rows**: In situ hybridization for **Neurogenin 2 (Ngn2)** at **E10.5** in control and **Dnm2 E-cKO** embryos. Compared to controls, **Dnm2 E-cKO** cortices display expanded and ectopic Ngn2 expression domains (arrows), consistent with premature or dysregulated neuronal differentiation. Images are representative of multiple embryos per genotype. Sections were processed and imaged under identical conditions.

**Figure S4.**
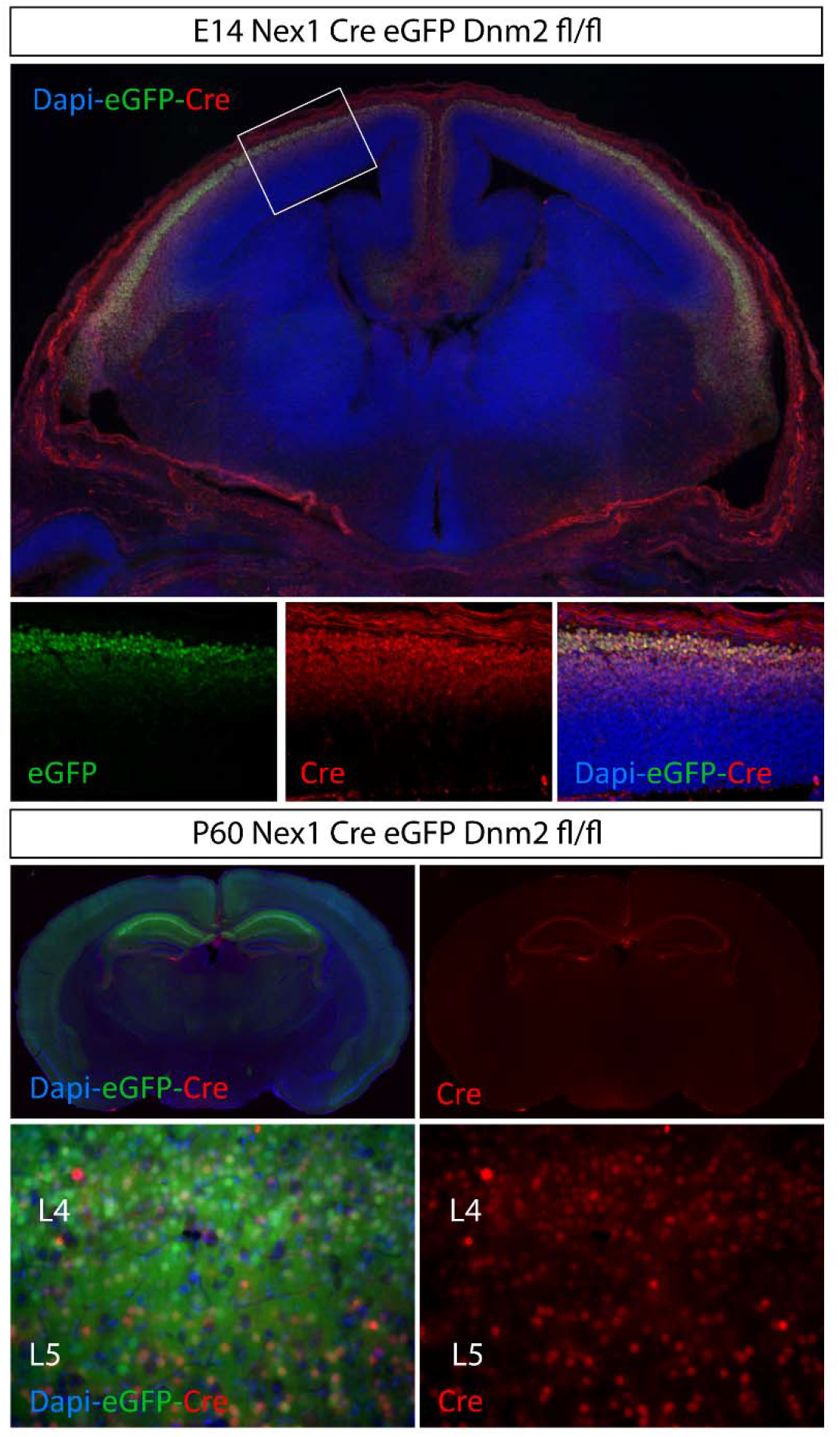
Nex1-Cre–mediated recombination in postmitotic cortical neurons. **Top panels**: Low-magnification coronal section of **E14 Nex1-Cre eGFP; Dnm2^fl/fl** cortex stained with **DAPI** (blue), **eGFP** (green), and **Cre** (red). Nex1-Cre–driven recombination is restricted to the dorsal cortical plate, with minimal signal in ventricular or subventricular progenitor zones. The boxed region indicates the area shown at higher magnification below. **Middle panels**: Higher-magnification views of the boxed cortical region showing robust overlap between **eGFP** and **Cre** expression in postmitotic cortical neurons, confirming efficient and spatially restricted recombination at E14. **Bottom panels**: Coronal sections from **P60 Nex1-Cre eGFP; Dnm2^fl/fl** brains stained for **DAPI, eGFP**, and **Cre**, demonstrating persistent Cre expression and recombination in mature cortex. Higher-magnification images highlight labeled neurons across cortical layers **L4** and **L5**, indicating long-term maintenance of Nex1-Cre–mediated recombination in postmitotic excitatory neurons. Images are representative of multiple animals. Sections were processed and imaged using identical conditions.

**Figure S5.**
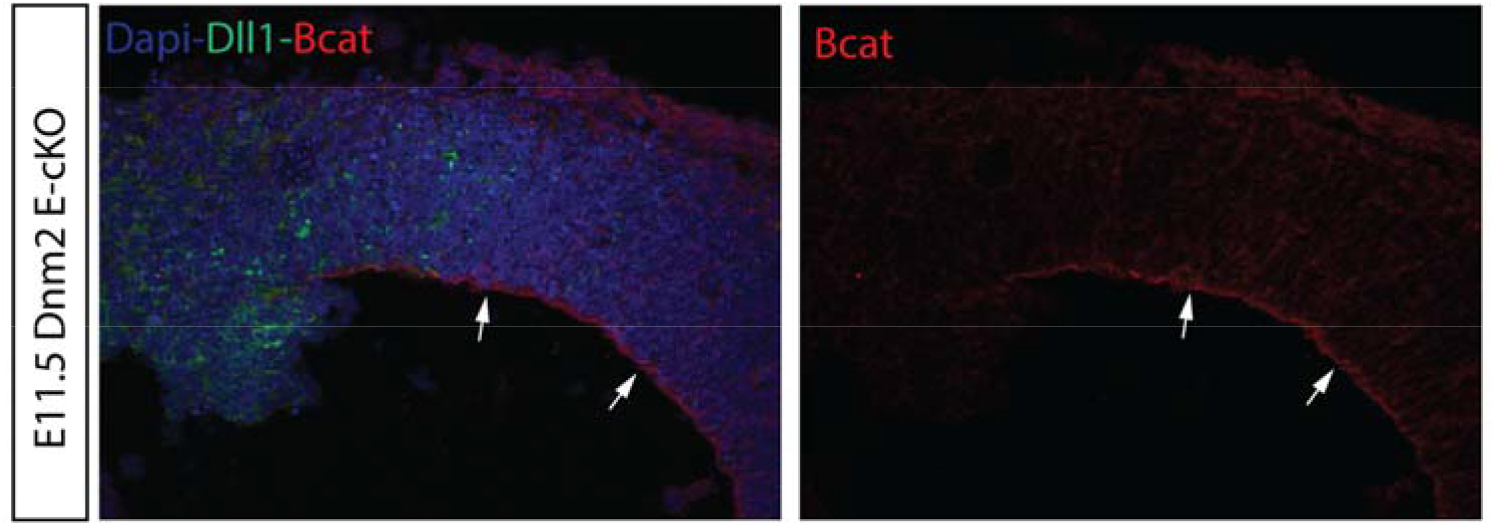
Altered β-catenin distribution in the ventricular zone following Dnm2 deletion. Immunofluorescence analysis **of E11.5 Dnm2 E-cKO** embryonic cortex stained with **DAPI** (blue), **Dll1** (green), and **β-catenin (Bcat)** (red). Left panel shows the merged image; right panel shows **β-catenin** signal alone. In **Dnm2 E-cKO** cortices, β-catenin signal is prominently enriched along the ventricular surface (arrows), coinciding with regions of altered ventricular zone architecture. Dll1-positive cells are distributed throughout the cortical wall.

**Figure S6.**
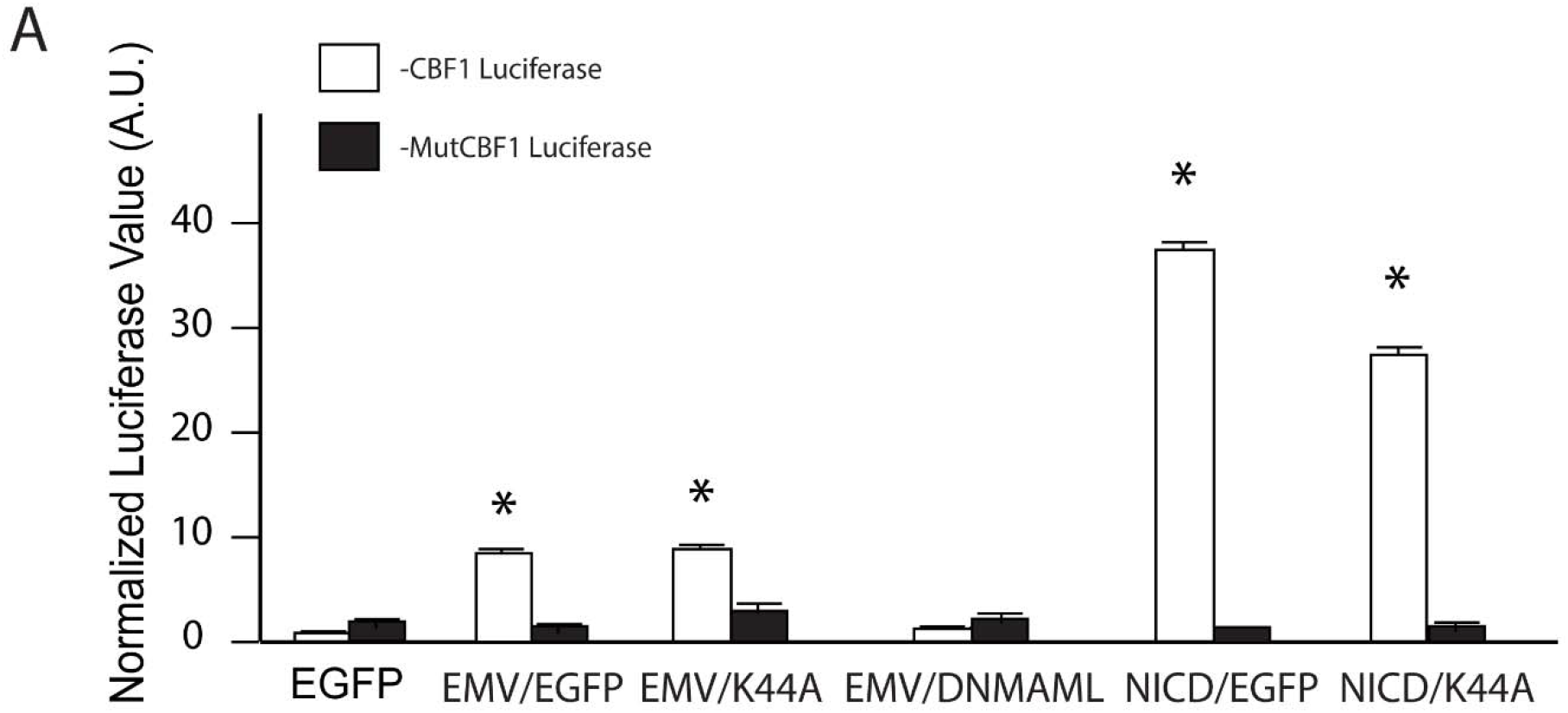
Dominant-negative Dynamin does not suppress signaling from EMV Notch. Neural stem cells were transfected with EMV Notch, a Notch construct lacking the extracellular domain and constitutively processed by γ-secretase, together with either EGFP or dominant-negative Dynamin 2 (K44A). Notch-dependent luciferase activity was quantified as in Figure 7. No reduction in reporter activity was observed following Dynamin inhibition. **(A)** Dual-luciferase reporter assay measuring Notch-dependent transcription using a **CBF1-luciferase reporter** (white bars) or a **mutated CBF1 reporter (MutCBF1)** (black bars). Cells were transfected with **EGFP**, empty vector (EMV) with EGFP, **Dynamin2 K44A, dominant-negative MAML (DNMAML), NICD**, or **NICD plus Dynamin2 K44A**, as indicated. Expression of **NICD** robustly activated CBF1-luciferase activity, whereas the MutCBF1 reporter remained at baseline, confirming reporter specificity. In contrast, expression of **Dynamin2 K44A** or **DNMAML** failed to activate CBF1-dependent transcription and suppressed Notch reporter activity relative to controls. Co-expression of NICD with Dynamin2 K44A significantly reduced **NICD**-driven transcriptional activation. Luciferase values were normalized to internal controls and are presented as mean ± SEM. Asterisks indicate statistical significance compared to corresponding controls (*p < 0.05).

## REFERENCES

Ables, J. L., J. J. Breunig, A. J. Eisch and P. Rakic (2011). “Not(ch) just development: Notch signalling in the adult brain.” Nat Rev Neurosci 12(5): 269–283.

Alberi, L., S. Liu, Y. Wang, R. Badie, C. Smith-Hicks, J. Wu, T. J. Pierfelice, B. Abazyan, M. P. Mattson, D. Kuhl, M. Pletnikov, P. F. Worley and N. Gaiano (2011). “Activity-induced Notch signaling in neurons requires Arc/Arg3.1 and is essential for synaptic plasticity in hippocampal networks.” Neuron 69(3): 437–444.

Andersson, E. R., R. Sandberg and U. Lendahl (2011). “Notch signaling: simplicity in design, versatility in function.” Development 138(17): 3593–3612.

Beckers, J., A. Caron, M. Hrabé de Angelis, S. Hans, J. A. Campos-Ortega and A. Gossler (2000). “Distinct regulatory elements direct delta1 expression in the nervous system and paraxial mesoderm of transgenic mice.” Mechanisms of development 95(1): 23–34.

Bray, S. J. (2006). “Notch signalling: a simple pathway becomes complex.” Nat Rev Mol Cell Biol 7(9): 678–689.

Breunig, J. J., T. F. Haydar and P. Rakic (2011). “Neural stem cells: historical perspective and future prospects.” Neuron 70(4): 614–625.

Breunig, J. J., R. Levy, C. D. Antonuk, J. Molina, M. Dutra-Clarke, H. Park, A. A. Akhtar, G. B. Kim, X. Hu, S. I. Bannykh, R. G. Verhaak and M. Danielpour (2015). “Ets Factors Regulate Neural Stem Cell Depletion and Gliogenesis in Ras Pathway Glioma.” Cell Rep 12(2): 258–271.

Breunig, J. J., J. Silbereis, F. M. Vaccarino, N. Sestan and P. Rakic (2007). “Notch regulates cell fate and dendrite morphology of newborn neurons in the postnatal dentate gyrus.” Proc Natl Acad Sci U S A 104(51): 20558–20563.

Bystron, I., C. Blakemore and P. Rakic (2008). “Development of the human cerebral cortex: Boulder Committee revisited.” Nat Rev Neurosci 9(2): 110–122.

Cao, H., F. Garcia and M. A. McNiven (1998). “Differential distribution of dynamin isoforms in mammalian cells.” Mol Biol Cell 9(9): 2595–2609.

Chou, S. J., C. G. Perez-Garcia, T. T. Kroll and D. D. O’Leary (2009). “Lhx2 specifies regional fate in Emx1 lineage of telencephalic progenitors generating cerebral cortex.” Nat Neurosci 12(11): 1381–1389.

Conner, S. D. and S. L. Schmid (2003). “Regulated portals of entry into the cell.” Nature 422(6927): 37–44.

Cook, T., K. Mesa and R. Urrutia (1996). “Three dynamin-encoding genes are differentially expressed in developing rat brain.” J Neurochem 67(3): 927–931.

Ferguson, S. M., G. Brasnjo, M. Hayashi, M. Wolfel, C. Collesi, S. Giovedi, A. Raimondi, L. W. Gong, P. Ariel, S. Paradise, E. O’Toole, R. Flavell, O. Cremona, G. Miesenbock, T. A. Ryan and P. De Camilli (2007). “A selective activity-dependent requirement for dynamin 1 in synaptic vesicle endocytosis.” Science 316(5824): 570–574.

Ferguson, S. M. and P. De Camilli (2012). “Dynamin, a membrane-remodelling GTPase.” Nat Rev Mol Cell Biol 13(2): 75–88.

Ferguson, S. M., A. Raimondi, S. Paradise, H. Shen, K. Mesaki, A. Ferguson, O. Destaing, G. Ko, J. Takasaki, O. Cremona, O. T. E and P. De Camilli (2009). “Coordinated actions of actin and BAR proteins upstream of dynamin at endocytic clathrin-coated pits.” Dev Cell 17(6): 811–822.

Fortini, M. E. and D. Bilder (2009). “Endocytic regulation of Notch signaling.” Curr Opin Genet Dev 19(4): 323–328.

Frederiksen, K. and R. D. McKay (1988). “Proliferation and differentiation of rat neuroepithelial precursor cells in vivo.” J Neurosci 8(4): 1144–1151.

Goebbels, S., I. Bormuth, U. Bode, O. Hermanson, M. H. Schwab and K. A. Nave (2006). “Genetic targeting of principal neurons in neocortex and hippocampus of NEX-Cre mice.” Genesis 44(12): 611–621.

Gupta-Rossi, N., E. Six, O. LeBail, F. Logeat, P. Chastagner, A. Olry, A. Israel and C. Brou (2004). “Monoubiquitination and endocytosis direct gamma-secretase cleavage of activated Notch receptor.” J Cell Biol 166(1): 73–83.

Guruharsha, K. G., M. W. Kankel and S. Artavanis-Tsakonas (2012). “The Notch signalling system: recent insights into the complexity of a conserved pathway.” Nat Rev Genet 13(9): 654–666.

Hockfield, S. and R. D. McKay (1985). “Identification of major cell classes in the developing mammalian nervous system.” J Neurosci 5(12): 3310–3328.

Itoh, M., C.-H. Kim, G. Palardy, T. Oda, Y.-J. Jiang, D. Maust, S.-Y. Yeo, K. Lorick, G. J. Wright and L. Ariza-McNaughton (2003). “Mind bomb is a ubiquitin ligase that is essential for efficient activation of Notch signaling by Delta.” Developmental cell 4(1): 67–82.

Iwasato, T., R. Nomura, R. Ando, T. Ikeda, M. Tanaka and S. Itohara (2004). “Dorsal telencephalon-specific expression of Cre recombinase in PAC transgenic mice.” Genesis 38(3): 130–138.

Jaekel, R. and T. Klein (2006). “The Drosophila Notch Inhibitor and Tumor Suppressor Gene lethal (2) giant discs Encodes a Conserved Regulator of Endosomal Trafficking.” Developmental cell 11(5): 655–669.

Kadowaki, M., S. Nakamura, O. Machon, S. Krauss, G. L. Radice and M. Takeichi (2007). “N-cadherin mediates cortical organization in the mouse brain.” Developmental Biology 304(1): 22–33.

Kanwar, R. and M. E. Fortini (2008). “The big brain aquaporin is required for endosome maturation and notch receptor trafficking.” Cell 133(5): 852–863.

Kawaguchi, A., T. Ikawa, T. Kasukawa, H. R. Ueda, K. Kurimoto, M. Saitou and F. Matsuzaki (2008). “Single-cell gene profiling defines differential progenitor subclasses in mammalian neurogenesis.” Development 135(18): 3113–3124.

Kawaguchi, D., T. Yoshimatsu, K. Hozumi and Y. Gotoh (2008). “Selection of differentiating cells by different levels of delta-like 1 among neural precursor cells in the developing mouse telencephalon.” Development 135(23): 3849–3858.

Kim, G. B., D. Rincon Fernandez Pacheco, D. Saxon, A. Yang, S. Sabet, M. Dutra-Clarke, R. Levy, A. Watkins, H. Park, A. Abbasi Akhtar, P. W. Linesch, N. Kobritz, S. S. Chandra, K. Grausam, A. Ayala-Sarmiento, J. Molina, K. Sedivakova, K. Hoang, J. Tsyporin, D. S. Gareau, M. G. Filbin, S. Bannykh, C. Santiskulvong, Y. Wang, J. Tang, M. L. Suva, B. Chen, M. Danielpour and J. J. Breunig (2019). “Rapid Generation of Somatic Mouse Mosaics with Locus-Specific, Stably Integrated Transgenic Elements.” Cell 179(1): 251–267 e224.

Koo, B. K., M. J. Yoon, K. J. Yoon, S. K. Im, Y. Y. Kim, C. H. Kim, P. G. Suh, Y. N. Jan and Y. Y. Kong (2007). “An obligatory role of mind bomb-1 in notch signaling of mammalian development.” PLoS One 2(11): e1221.

Kopan, R. and M. X. Ilagan (2009). “The canonical Notch signaling pathway: unfolding the activation mechanism.” Cell 137(2): 216–233.

Kriegstein, A. and A. Alvarez-Buylla (2009). “The glial nature of embryonic and adult neural stem cells.” Annu Rev Neurosci 32: 149–184.

Lai, E. C. and G. M. Rubin (2001). “Neuralized is essential for a subset of Notch pathway-dependent cell fate decisions during Drosophila eye development.” Proceedings of the National Academy of Sciences 98(10): 5637–5642.

Maillard, I., A. P. Weng, A. C. Carpenter, C. G. Rodriguez, H. Sai, L. Xu, D. Allman, J. C. Aster and W. S. Pear (2004). “Mastermind critically regulates Notch-mediated lymphoid cell fate decisions.” Blood 104(6): 1696–1702.

Meloty-Kapella, L., B. Shergill, J. Kuon, E. Botvinick and G. Weinmaster (2012). “Notch ligand endocytosis generates mechanical pulling force dependent on dynamin, epsins, and actin.” Dev Cell 22(6): 1299–1312.

Mizutani, K., K. Yoon, L. Dang, A. Tokunaga and N. Gaiano (2007). “Differential Notch signalling distinguishes neural stem cells from intermediate progenitors.” Nature 449(7160): 351–355.

Nelson, B. R., R. D. Hodge, F. Bedogni and R. F. Hevner (2013). “Dynamic interactions between intermediate neurogenic progenitors and radial glia in embryonic mouse neocortex: potential role in Dll1-Notch signaling.” J Neurosci 33(21): 9122–9139.

Noctor, S. C., V. Martinez-Cerdeno, L. Ivic and A. R. Kriegstein (2004). “Cortical neurons arise in symmetric and asymmetric division zones and migrate through specific phases.” Nat Neurosci 7(2): 136–144.

Ohtsuka, T., I. Imayoshi, H. Shimojo, E. Nishi, R. Kageyama and S. K. McConnell (2006). “Visualization of embryonic neural stem cells using Hes promoters in transgenic mice.” Molecular and Cellular Neuroscience 31(1): 109–122.

Park, R. J., H. Shen, L. Liu, X. Liu, S. M. Ferguson and P. De Camilli (2013). “Dynamin triple knockout cells reveal off target effects of commonly used dynamin inhibitors.” J Cell Sci 126(Pt 22): 5305–5312.

Pasternak, S. H., R. D. Bagshaw, M. Guiral, S. Zhang, C. A. Ackerley, B. J. Pak, J. W. Callahan and D. J. Mahuran (2003). “Presenilin-1, nicastrin, amyloid precursor protein, and gamma-secretase activity are co-localized in the lysosomal membrane.” J Biol Chem 278(29): 26687–26694.

Qiu, C., B. K. Martin, I. C. Welsh, R. M. Daza, T.-M. Le, X. Huang, E. K. Nichols, M. L. Taylor, O. Fulton, D. R. O’Day, A. R. Gomes, S. Ilcisin, S. Srivatsan, X. Deng, C. M. Disteche, W. S. Noble, N. Hamazaki, C. B. Moens, D. Kimelman, J. Cao, A. F. Schier, M. Spielmann, S. A. Murray, C. Trapnell and J. Shendure (2024). “A single-cell time-lapse of mouse prenatal development from gastrula to birth.” Nature 626(8001): 1084–1093.

Raimondi, A., S. M. Ferguson, X. Lou, M. Armbruster, S. Paradise, S. Giovedi, M. Messa, N. Kono, J. Takasaki, V. Cappello, E. O’Toole, T. A. Ryan and P. De Camilli (2011). “Overlapping role of dynamin isoforms in synaptic vesicle endocytosis.” Neuron 70(6): 1100–1114.

Rakic, P. (1972). “Mode of cell migration to the superficial layers of fetal monkey neocortex.” J Comp Neurol 145(1): 61–83.

Rasin, M. R., V. R. Gazula, J. J. Breunig, K. Y. Kwan, M. B. Johnson, S. Liu-Chen, H. S. Li, L. Y. Jan, Y. N. Jan, P. Rakic and N. Sestan (2007). “Numb and Numbl are required for maintenance of cadherin-based adhesion and polarity of neural progenitors.” Nat Neurosci 10(7): 819–827.

Ring, K. L., L. M. Tong, M. E. Balestra, R. Javier, Y. Andrews-Zwilling, G. Li, D. Walker, W. R. Zhang, A. C. Kreitzer and Y. Huang (2012). “Direct reprogramming of mouse and human fibroblasts into multipotent neural stem cells with a single factor.” Cell Stem Cell 11(1): 100–109.

Sestan, N., S. Artavanis-Tsakonas and P. Rakic (1999). “Contact-dependent inhibition of cortical neurite growth mediated by notch signaling.” Science 286(5440): 741–746.

Seugnet, L., P. Simpson and M. Haenlin (1997). “Requirement for dynamin during Notch signaling in Drosophila neurogenesis.” Dev Biol 192(2): 585–598.

Shaye, D. D. and I. Greenwald (2002). “Endocytosis-mediated downregulation of LIN-12/Notch upon Ras activation in Caenorhabditis elegans.” Nature 420(6916): 686–690.

Simeone, A., D. Acampora, M. Gulisano, A. Stornaiuolo and E. Boncinelli (1992). “Nested expression domains of four homeobox genes in developing rostral brain.” Nature 358(6388): 687–690.

Sorensen, E. B. and S. D. Conner (2010). “gamma-secretase-dependent cleavage initiates notch signaling from the plasma membrane.” Traffic 11(9): 1234–1245.

Struhl, G. and A. Adachi (2000). “Requirements for presenilin-dependent cleavage of notch and other transmembrane proteins.” Mol Cell 6(3): 625–636.

Tagami, S., M. Okochi, K. Yanagida, A. Ikuta, A. Fukumori, N. Matsumoto, Y. Ishizuka-Katsura, T. Nakayama, N. Itoh, J. Jiang, K. Nishitomi, K. Kamino, T. Morihara, R. Hashimoto, T. Tanaka, T. Kudo, S. Chiba and M. Takeda (2008). “Regulation of Notch signaling by dynamic changes in the precision of S3 cleavage of Notch-1.” Mol Cell Biol 28(1): 165–176.

Tronche, F., C. Kellendonk, O. Kretz, P. Gass, K. Anlag, P. C. Orban, R. Bock, R. Klein and G. Schutz (1999). “Disruption of the glucocorticoid receptor gene in the nervous system results in reduced anxiety.” Nat Genet 23(1): 99–103.

Vaccari, T., H. Lu, R. Kanwar, M. E. Fortini and D. Bilder (2008). “Endosomal entry regulates Notch receptor activation in Drosophila melanogaster.” J Cell Biol 180(4): 755–762.

Windler, S. L. and D. Bilder (2010). “Endocytic internalization routes required for delta/notch signaling.” Curr Biol 20(6): 538–543.

Yoon, K. and N. Gaiano (2005). “Notch signaling in the mammalian central nervous system: insights from mouse mutants.” Nat Neurosci 8(6): 709–715.

Yoon, K. J., B. K. Koo, S. K. Im, H. W. Jeong, J. Ghim, M. C. Kwon, J. S. Moon, T. Miyata and Y. Y. Kong (2008). “Mind bomb 1-expressing intermediate progenitors generate notch signaling to maintain radial glial cells.” Neuron 58(4): 519–531.

Yoshida, M., Y. Suda, I. Matsuo, N. Miyamoto, N. Takeda, S. Kuratani and S. Aizawa (1997). “Emx1 and Emx2 functions in development of dorsal telencephalon.” Development 124(1): 101–111.

